# Spatiotemporal regulation of *de novo* and salvage purine synthesis during brain development

**DOI:** 10.1101/2023.03.01.530588

**Authors:** Tomoya Mizukoshi, Seiya Yamada, Shin-ichi Sakakibara

**Author notes:** To whom correspondence should be addressed: Shin-ichi Sakakibara or Seiya Yamada, Laboratory for Molecular Neurobiology, Faculty of Human Sciences, Waseda University, Tokorozawa, Saitama 359-1192, Japan; or. Funding This work was funded by the Japan Society for the Promotion of Science grants-in-aid (KAKENHI) grant numbers 21K20701 (to S.Y.) and 19K06931 (to S.S.), Gout and Uric Acid Research Foundation 2020 (to S. S) and 2022 (to S.Y.), and Waseda University Grants for Special Research Projects 2021C-611 (to S.Y.).

## Abstract

The levels of purines, essential molecules to sustain eukaryotic cell homeostasis, are regulated by the coordination of the *de novo* and salvage synthesis pathways. In the embryonic central nervous system (CNS), the *de novo* pathway is considered crucial to meet the requirements for the active proliferation of neural stem/progenitor cells (NSPCs). However, how these two pathways are balanced or separately utilized during CNS development remains poorly understood. In this study, we showed a dynamic shift in pathway utilization, with greater reliance on the *de novo* pathway during embryonic stages and on the salvage pathway at postnatal–adult stages. The pharmacological effects of various purine synthesis inhibitors *in vitro* and the expression profile of purine synthesis enzymes indicated that NSPCs in the embryonic cerebrum mainly utilize the *de novo* pathway. Simultaneously, NSPCs in the cerebellum require both the *de novo* and the salvage pathways. *In vivo* administration of *de novo* inhibitors resulted in severe hypoplasia of the forebrain cortical region, indicating a gradient of purine demand along the anteroposterior axis of the embryonic brain, with cortical areas of the dorsal forebrain having higher purine requirements than ventral or posterior areas such as the striatum and thalamus. This histological defect of the neocortex was accompanied by strong downregulation of the mechanistic target of rapamycin complex 1 (mTORC1)/ribosomal protein S6 kinase (S6K)/S6 signaling cascade, a crucial pathway for cell metabolism, growth, and survival. These findings indicate the importance of the spatiotemporal regulation of both purine pathways for mTORC1 signaling and proper brain development.

**Significance Statement:** Brain development requires a balance of *de novo* and salvage purine synthetic pathways. However, the utilization of these pathways during brain development remains poorly understood. This study provides evidence that the spatiotemporal regulation of these two purine synthesis pathways is essential for normal brain development. We revealed that inhibition of *de novo* purine synthesis results in the downregulation of mammalian/mechanistic target of rapamycin (mTOR) signaling, leading to malformations in specific embryonic brain regions such as the cerebral neocortex. These results suggest a temporal and spatial gradient of purine demand during embryonic brain development. These findings could improve our understanding of neurological diseases caused by defects in purine metabolism.

## Introduction

The spatiotemporally regulated proliferation of neural stem cells and the migration of newborn neurons are crucial for forming the mammalian cerebral cortex. In the embryonic mouse brain, neural stem cells actively and symmetrically proliferate in the ventricular zone (VZ) to expand their original pool, eventually dividing asymmetrically to generate either intermediate progenitor cells (IPs) or neurons. Here, the neural stem cells in the VZ and the IPs in the subventricular zone (SVZ) are collectively referred to as neural stem/progenitor cells (NSPCs). Numerous glutamatergic neurons generated from NSPCs migrate radially to form the stratified layers of the cerebral cortex (Agirman et al., 2017). In addition, many GABAergic inhibitory neurons, generated in the embryonic ganglionic eminence (GE), tangentially migrate, finally integrating in the cerebral cortex (Lavdas et al., 1999; Cho et al., 2014; Corbin et al., 2003). Defects in radial migration frequently cause brain malformation and psychiatric disorders, including mental retardation and epilepsy (Moffat et al., 2015).

Purines are the essential building blocks of DNA/RNA, energy sources of enzymatic reactions (ATP/GTP), and second messengers in various intracellular signaling cascades (cyclic AMP and cyclic GMP). Moreover, adenosine and ATP are involved in extracellular signaling mediated by purinergic receptors, regulating cell functions, such as cell migration, apoptosis, NSPC proliferation, and neuron differentiation (Huang et al., 2021; Lin et al., 2007).

There are two pathways for purine synthesis in mammals: the *de novo* and salvage pathways. In the latter, purines are synthesized by recycling degraded bases (AMP, GMP, and IMP) derived from the catabolism (Jurecka, 2009). The hypoxanthine-guanine phosphoribosyl transferase (HGPRT) converts hypoxanthine and guanine into inosine monophosphate (IMP) and guanine monophosphate (GMP), respectively, whereas the adenine phosphoribosyltransferase synthesizes AMP from adenine (Fig. 1) (Pedley and Benkovic, 2017). Under physiological conditions, purines are generally supplied primarily via the salvage pathway. The *de novo* pathway is supposed to be set off when the demand for purines exceeds the capacity of salvage synthesis (Pareek et al., 2020). The *de novo* pathway produces IMP from phosphoribosyl pyrophosphate through 10 consecutive reactions mediated by six enzymes [phosphoribosyl pyrophosphate amidotransferase, phosphoribosylglycinamide formyltransferase, formylglycinamidine ribonucleotide synthase (FGAMS), phosphoribosylaminoimidazolecarboxylase phosphoribosyl-aminoimidazole succinocarboxamide synthetase (PAICS), adenylosuccinate lyase (ADSL), and 5-aminoimidazole-4-carboxamide ribonucleotide formyltransferase/IMP cyclohydrolase (ATIC)] (Fig. 1) (Pareek et al., 2021). These six *de novo* enzymes are assembled into a huge complex called purinosome, enabling a higher efficacy of IMP synthesis (An et al., 2008; French et al., 2016; Pedley and Benkovic, 2017). In fact, we recently demonstrated that a defect in purinosome formation leads to malformation of the cerebral cortex (Yamada et al., 2020). Other studies also provided evidence regarding a close relationship between purine production and the mammalian/mechanistic target of the rapamycin complex 1 (mTORC1) signaling pathway (Ben-Sahra et al., 2016; Hoxhaj et al., 2017); however, the molecular interplay between mTORC1 signaling and purine metabolism in cortical development remains unclear.

**Figure 1.**
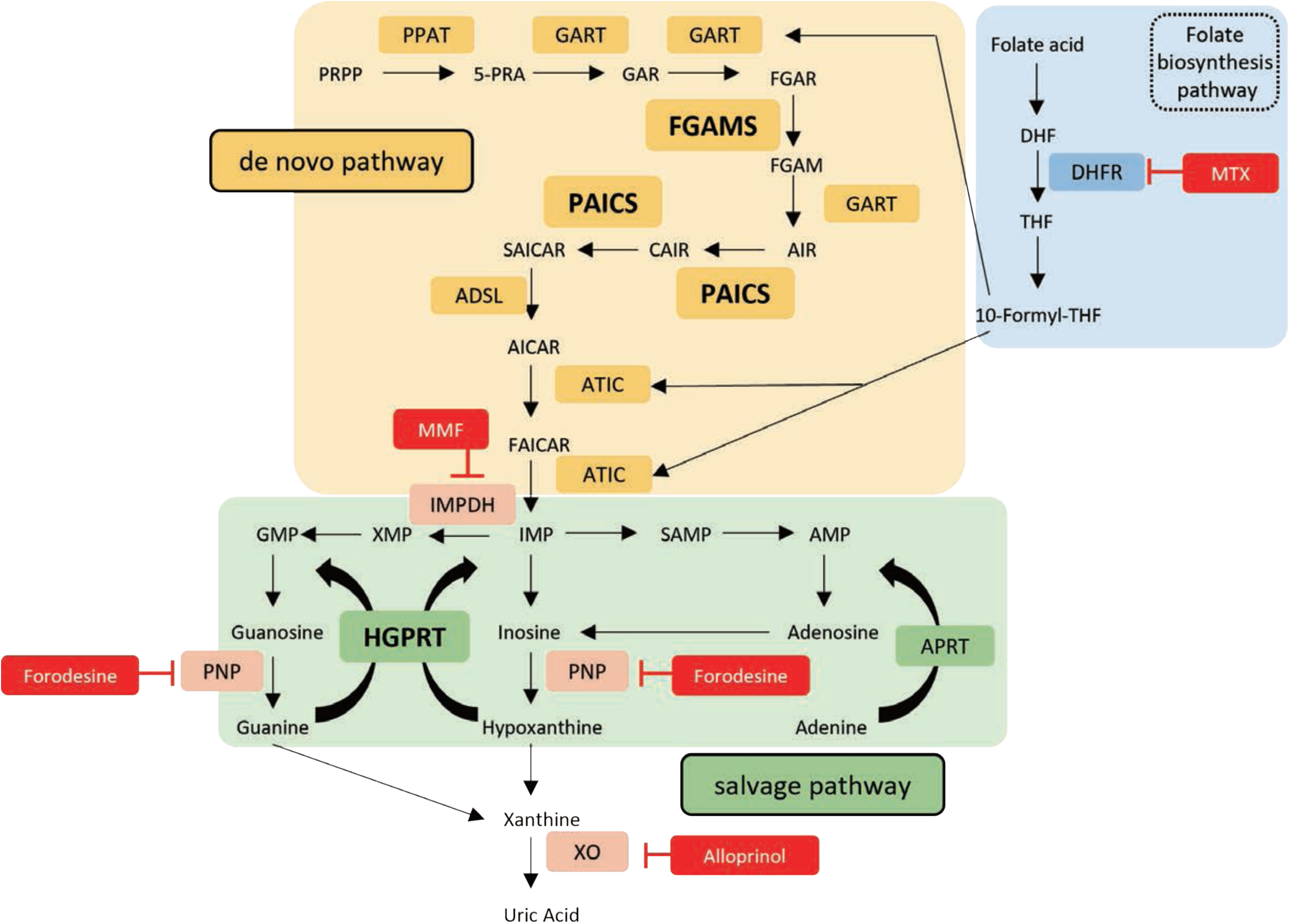
*De novo* and salvage purine synthesis pathways in mammals. Metabolic diagram of *de novo* (PAICS, FGAMS) and salvage (HGPRT) purine synthesis pathways in mammals. Enzymes involved in the *de novo* and salvage pathways are represented by yellow and green boxes, respectively. The route in blue represents the folate biosynthesis pathway, which is essential for substrate production of *de novo* purine synthesis. Red boxes indicate the inhibitors used in this study. PRPP, 5-phosphoribosyl-1-pyrophosphate; PPAT, phosphoribosyl pyrophosphate amidotransferase; 5-PRA, 5-phosphoribosylamine; GART, phosphoribosylglycinamide formyltransferase; GAR, glycinamide ribonucleotide; FGAR, formylglycinamide ribonucleotide; FGAMS, formylglycinamidine ribonucleotide synthase; FGAM, formylglycinamidine ribonucleotide; AIR, aminoimidazole ribonucleotide; PAICS, phosphoribosylaminoimidazole succinocarboxamide synthetase; CAIR, 4-carboxy-5-aminoimidazole ribonucleotide; SAICAR, 4-(N-succinylcarboxamide)-5-aminoimidazole ribonucleotide; ADSL, adenylosuccinate lyase; AICAR, aminoimidazole-4-carboxamide ribonucleotide; FAICAR, formylaminoimidazole-4-carboxamide ribonucleotide; ATIC, 5-aminoimidazole-4-carboxamide ribonucleotide formyltransferase inosine monophosphate cyclohydrolase; DHF, dihydrofolate; DHFR, dihydrofolate reductase; THF, tetrahydrofolic acid; HGPRT, hypoxanthine-guanine phosphoribosyltransferase; APRT, adenine phosphoribosyltransferase; IMP, inosine monophosphate; IMPDH, inosine monophosphate dehydrogenase; XMP, xanthosine monophosphate; GMP, guanosine monophosphate; SAMP, adenylosuccinate; AMP, adenosine monophosphate; PNP, purine nucleoside phosphorylase; XO, xanthine oxidase; MMF, mycophenolate mofetil; MTX, methotrexate; Forodesine, forodesine hydrochloride.

Furthermore, abnormalities in the purine metabolism are associated with the etiology of many diseases, including gout. For instance, inherited deficiencies in *de novo* enzymes result in fetal lethality or neurological diseases in humans (Jurecka, 2009). For instance, a missense mutation in *PAICS* causes multiple malformations and early neonatal death (Pelet et al., 2019), and *ADSL* or *ATIC* deficiency causes various neurodevelopmental phenotypes, including epilepsy, speech impairment, and auto-aggressive behavior (Marie et al., 2004; Jurecka et al., 2015; Dutto et al., 2022). With respect to the salvage pathway, *HGPRT* deficiency causes the Lesch-Nyhan syndrome, characterized by juvenile gout, dystonia, mental retardation, and compulsive self-injurious behavior (Lesch and Nyhan, 1964; Torres and Puig, 2007). Although *HGPRT* deficiency is thought to cause functional impairment of the dopamine system in the basal ganglia (Visser et al., 2000), no fundamental therapeutic approach has been established.

These previous studies strongly suggest that the balance between *de novo* and salvage purine synthesis pathways is critical for healthy brain development. However, the spatiotemporal regulation of purine production pathways in CNS development remains unknown. Here, we report the expression profile and the functional significance of purine synthesis enzymes in the developing brain. Each purine pathway is driven by its own temporal and regional traits during brain development.

## Materials and Methods

### Animals

ICR and C57BL/6J mice (Japan SLC Inc.) were housed under temperature- and humidity-controlled conditions under a 12/12-h light/dark cycle, with *ad libitum* access to food and water. The date of conception was established using a vaginal plug and recorded as embryonic day zero (E0.5). The day of birth was designated as P0.

### Tissue preparation

Embryonic and adult mice were perfused through the cardiac ventricle with 4% paraformaldehyde (PFA) in 0.1 M phosphate buffer (pH 7.4), followed by post-fixation overnight at 4°C (Yamada and Sakakibara, 2018). Fixed brains were cryoprotected in 30% sucrose in phosphate-buffered saline (PBS) overnight at 4℃ and embedded in the optimal cutting temperature compound (Sakura Finetek). Frozen sections of embryonic and postnatal brains were cut into 14 μm thickness sections using a cryostat and collected on MAS-coated glass slides (Matsunami Glass). Free-floating sections of adult mice brains were cut at 30 μm thickness.

### Primary cell culture

NSPCs were cultured as described previously (Yamada and Sakakibara, 2018; Yumoto et al., 2013). They were first isolated from the E12.5 telencephalon or P2 external granular layer of the cerebellum, seeded onto dishes coated with 5 µg/mL fibronectin and 0.2% polyethyleneimine (Merck Millipore), and cultured in Advanced DMEM/F-12 (1:1) (Thermo Fisher Scientific) supplemented with 15 μg/mL insulin (Thermo Fisher Scientific), 25 μg/mL transferrin (Thermo Fisher Scientific), 20 nM progesterone (Merck Millipore), 30 nM sodium selenite (Merck Millipore), 60 nM putrescine (Merck Millipore), 1% penicillin-streptomycin (Fujifilm Wako), 20 ng/mL fibroblast growth factor 2 (Merck Millipore), and 10 ng/mL epidermal growth factor (Merck Millipore). To assess cell proliferation, 10 μM 5-bromodeoxyuridine (BrdU) (Tokyo Chemical Industry) was administered to the cultured NSPCs. After 24 h of incubation, cells were cultivated in the medium without BrdU for the indicated periods and processed for immunostaining using an anti-BrdU antibody. Electroporation of Paics shRNAs into NSPCs was performed as described previously (Yamada et al., 2020). The validated targeting sequences for Paics shRNA #1 and shRNA #3 were 5′-CTGCTCAGATATTTGGGTTAA-3′ and 5′-GCACCTGCTTTCAAATACTAT-3′, respectively (Yamada et al., 2020). The NSPCs expanded *in vitro* were electroporated with Paics shRNAs using a NEPA21 electroporator (Nepagene) according to the manufacturer’s specifications (NSPCs: two pulses of 125 V for 5 ms with a 50 ms interval). After 96 h of culturing, cells were incubated with 10 μM BrdU for 24 h and fixed. To obtain primary cultured neurons, E16.5 cerebral cortices or P4 cerebellar cortices were dissected and dispersed as previously described (Yamada et al., 2021; Yamada et al., 2022). Cells were seeded onto poly-D-lysine-coated dishes (Merck Millipore) and cultured in a neurobasal medium containing 2% B27 (Thermo Fisher Scientific) and 1% GlutaMax (Thermo Fisher Scientific) for 10 days *in vitro* (div). Primary cultured astrocytes differentiated from NSPCs prepared from E12.5 cerebral cortices or P2 cerebella after being cultured for five divisions in DMEM (Fujifilm Wako) supplemented with 10% fetal bovine serum (Thermo Fisher Scientific), 1% L-glutamine (Fujifilm Wako), and 1% penicillin–streptomycin.

### Immunohistochemistry

Immunohistochemistry (IHC) was performed as described previously (Yamada and Sakakibara, 2018). For antigen retrieval, frozen sections were heated at 90°C–95℃ for 10 min in 10 mM sodium citrate buffer (pH 6.0) using a microwave and treated with 1% H_2_O_2_ in PBST (0.1% Triton X-100 in PBS) for 30 min at 20°C–25℃ to suppress the endogenous peroxidase activity. Sections were blocked for 2 h in 10% normal goat serum in PBST and incubated overnight at 4℃ with the primary antibodies in the blocking buffer. Then, the sections were incubated with biotinylated anti-rabbit IgG (Vectastain ABC Elite; Vector Lab) in PBST at room temperature for 2 h, followed by incubation with the ABC reagent (Vectastain ABC Elite; Vector Lab) in PBST for 2 h. The horseradish peroxidase signal was visualized using 2.25% diaminobenzidine and 2.25% H_2_O_2_ (Peroxidase Stain DAB Kit; Nacalai tesque). Each step was followed by four washes in PBST. Free-floating sections were mounted on MAS-coated glass slides, dehydrated, and coverslipped with Entellan New (Merck Millipore).

Double immunostaining was performed as described previously (Yamada and Sakakibara, 2018; Yamada et al., 2020). Frozen sections were blocked for 2 h with 5% normal goat or donkey serum in PBST, followed by incubation with the primary antibodies in blocking buffer at 4℃ overnight. After washing with PBST four times and twice with PBS, sections were incubated for 2 h with Alexa Fluor 488- or Alexa Fluor 555-conjugated secondary antibodies (Thermo Fisher Scientific). After counterstaining with 0.7 nM Hoechst 33342 (Thermo Fisher Scientific), sections were mounted and imaged using a confocal (FV3000, Olympus) or fluorescence inverted microscope (Axio Observer, Zeiss). For BrdU staining, sections were treated with 2 N HCl for 30 min at 42℃ for DNA denaturation, followed by treatment with 0.1 M sodium borate buffer (pH 8.5) for 10 min. After washing with PBST four times and twice with PBS, anti-BrdU immunostaining was performed.

### Immunocytochemistry

Cultured cells were fixed with 4% PFA and permeabilized with 0.1% Triton X-100. For BrdU staining, cells were treated with 2 N HCl for 1 h at 37℃, followed by neutralization with 0.1 M sodium borate buffer (pH 8.5) for 30 min. After blocking with 5% normal donkey serum, cells were incubated with primary antibody for 1 h at room temperature, followed by Alexa Fluor 488- or 594-conjugated secondary antibodies (Thermo Fisher Scientific) for 2 h. Immunofluorescent signals were acquired using an inverted microscope (Axio Observer, Zeiss) equipped with a CCD camera (AxioCam MRm, Zeiss).

### Primary antibodies

The following antibodies were used: anti-α-Tubulin (rabbit polyclonal, #11224-1-AP, RRID: AB_2210206, Proteintech; 1:5000 for Western blot, WB), anti-PAICS (rabbit polyclonal, #12967-1-AP, RRID: AB_10638449, Proteintech; 1:1500 for WB, 1:200 for IHC), anti-PFAS (FGAMS) (rabbit polyclonal, #A304-218A, RRID: AB_2620415, Bethyl Lab; 1:1500 for WB, 1:200 for IHC), anti-HPRT1 (HGRPT) (rabbit polyclonal, #15059-1-AP, RRID: AB_10638622, Proteintech; 1:3000 for WB, 1:1000 for IHC), anti-Pax6 (rabbit polyclonal, #PD022, RRID: AB_1520876, MBL; 1:3000 for WB), anti-GFAP (mouse monoclonal, #G3893, RRID: AB_477010, Sigma; 1:2000 for WB, 1:1000 for ICC), anti-CD31 (rat monoclonal clone ER-MP12, #MCA2388T, RRID: AB_1101899, BioRad Laboratory; 1:200 for IHC), anti-Nestin (chicken polyclonal, #GTX85457, RRID: AB_10627357, Gene Tex; 1:1500 for ICC), anti-BrdU (sheep polyclonal, #ab1893, RRID: AB_302659, Abcam, 1:1500 for IHC and ICC), anti-Ki67 (rabbit monoclonal clone SP6, #RM-9106, RRID: AB_2341197, Lab Vision; 1:1000 for IHC), anti-Phospho-Histone H3 (Ser10) (mouse monoclonal clone 6G3, #9706, RRID: AB_331748, Cell Signaling Technology; 1:1000 for IHC), anti-Phospho-p70 ribosomal protein S6 kinase (Thr389) (108D2) (pS6K) (rabbit monoclonal, #9234, RRID: AB_2269803, Cell Signaling Technology; 1:1000 for WB), anti-p70 S6K α (S6K)(mouse monoclonal clone H-9, #sc-8418, RRID: AB_628094, Santa Cruz Biotechnology; 1:1000 for WB), anti-Phospho-S6 ribosomal protein (Ser240/244) (pS6) (rabbit polyclonal, #2215, RRID: AB_331682, Cell Signaling Technology; 1:1000 for WB), anti-S6 ribosomal protein (S6) (mouse monoclonal clone C-8, #sc-74459, RRID: AB_1129205, Santa Cruz Biotechnology; 1:1000 for WB), anti-4E-BP1 (rabbit monoclonal clone 53H11, #9644, RRID: AB_2097841, Cell Signaling Technology; 1:1000 for WB), anti-cleaved-caspase3 (rabbit polyclonal, #9661S, RRID: AB_2910623, Cell Signaling; 1:500 for WB, ICC and IHC) and anti-GSH2 (rabbit polyclonal, #ABN162, RRID: AB_11203296, Sigma Aldrich; 1:200 for IHC).

### Drugs

The following drugs were used: mycophenolate mofetil (MMF) (#S1501, Selleck, 100 mg/kg *in vivo*, 1–10 μM *in vitro*, dissolved in DMSO), methotrexate (MTX) (#139-13571, Fujifilm Wako, 50 mg/kg *in vivo*, 1–100 μM *in vitro*, dissolved in 50 mM Na_2_CO_3_ buffer), forodesine hydrochloride (#HY-16209, MCE, 40 mg/kg *in vivo*, 0.1–5 μM *in vitro*, dissolved in PBS), allopurinol (#17795-21-0, Selleck, 50 mg/kg *in vivo*, 250 μM *in vitro*, dissolved in DMSO), BrdU (#B1575, Tokyo Chemical Industry, 100 mg/kg *in vivo*, 10 μM *in vitro*, dissolved in PBS), and MHY1485 (#S7811, Selleck, 50 mg/kg *in vivo*, 10 μM *in vitro*, dissolved in DMSO). For *in vitro* culture, drugs were added to the medium at the indicated concentration and incubated for 48 h. BrdU was added 24 h prior to fixation to mark S-phase cells. For *in vivo* experiments, the inhibitor was administered to pregnant dams or postnatal pups by a single intraperitoneal injection (i.p.) at E12.5, E16.5, or P2, followed by BrdU injection at E13.5, E17.5, or P4, respectively. Embryos or pups were fixed at E14.5, E18.5, or P6 in each condition. In some experiments, the inhibitor was injected daily into pregnant mice from E9.5 to E11.5, and the fetuses were fixed at E12.5.

### Western blotting

Tissue lysates were prepared by homogenization in RIPA buffer [25 mM Tris–HCl (pH 7.6), 150 mM NaCl, 1% NP40, 1% sodium deoxycholate, 0.1% sodium dodecyl sulfate, containing protease inhibitors (cOmplete Mini Protease Inhibitor Cocktail, Sigma Aldrich)], followed by centrifugation at 15,000 rpm for 10 min. Western blotting was performed as described previously (Yamada et al., 2020). For pS6 and pS6K detection, the membranes were blocked using 1% bovine serum albumin (BSA) fraction Ⅴ (Fujifilm Wako) in 0.1% Tween 20 in Tris-buffered saline (TBST), followed by incubation with a primary antibody diluted by 1% BSA in TBST. After washing with TBST, the membranes were incubated with horseradish peroxidase (HRP)-conjugated secondary antibody (Cytiva or Jackson ImmunoResearch). The signal was detected using Immobilon Western chemiluminescent HRP substrate (Merck Millipore). Blot images were captured using the Fusion Solo S system (Vilber Lourmat) and quantified using the Quantification Software Module (Vilber Lourmat).

### Statistical analyses

Statistical analyses were performed using the R package version 4.2.0. All numerical data are expressed as mean ± SEM. A one-way ANOVA followed by Weltch’s t-test with Holm–Bonferroni correction was used in multiple-group comparisons. Welch’s t-test was used to assess the number of BrdU^+^, pH3^+^, or cleaved-caspase3^+^ cells *in vivo* and pS6/S6 ratio. In *in vivo* experiments involving inhibitors, three individuals were tested per condition, and three images from each individual were randomly selected for statistical processing. The chi-squared test was used to compare the number of BrdU^+^ Nestin^+^, BrdU^+^ GFP^+^, or cleaved-caspase3^+^ Hoechst^+^ cells *in vitro*. *, P < 0.05; **, P < 0.01; ***, P <0.001 were considered statistically significant. The number of BrdU^+^ NSPCs *in vitro* was counted using MATLAB [version 9.12.0.1884302 (R2022a)]. The detailed statistical analyses are shown in Table 1.

## Results

### Two purine synthesis pathways are activated during brain development

To investigate the expression of purine synthesis enzymes in the developing CNS, immunoblotting analysis was performed on embryonic, postnatal, and adult whole brains. PAICS and FGAMS, both catalyzing the *de novo* purine synthesis (Fig. 1), were abundant in the embryonic stages from E10.5 to E16.5 during the active NSPC proliferation. Subsequently, from the postnatal to the adult stage, the expression of these *de novo* enzymes was downregulated (Fig. 2A). Conversely, the expression of HGPRT, a key enzyme catalyzing the salvage pathway (Fig. 1), was relatively low in embryonic brains, gradually increasing in postnatal and adult stages (Fig. 2A). Although HGPRT was expressed as a 25 kDa product with the expected size throughout life, an additional product of 35 kDa was detected during the embryonic and neonatal (E13–P0) period. Whether this slower-migrating band reflects another isoform or a post-translationally modified HGPRT product is unclear. To examine the expression properties of each brain region, we further divided the brain into the cerebral cortex (E13.5–P12) and cerebellum (P2–P12) and then performed the same experiments (Fig. 2B, C). The expression patterns of PAICS and HGPRT switched around the time of birth (P0) in the developing cerebral cortex; i.e., high PAICS expression was observed in early embryonic stages, whereas HGPRT expression increased after the postnatal period (Fig. 2B, C). Conversely, both PAICS and HGPRT were expressed at high levels in the P2 and P5 cerebellum, in which NSPCs actively divide to produce granule cells (Fig. 2B, C). Subsequently, HGPRT expression continued to increase in the cerebellum until P12 (Fig. 2B, C). These findings indicated that the choice of purine pathway depends on the stage of development and region of the brain.

**Figure 2.**
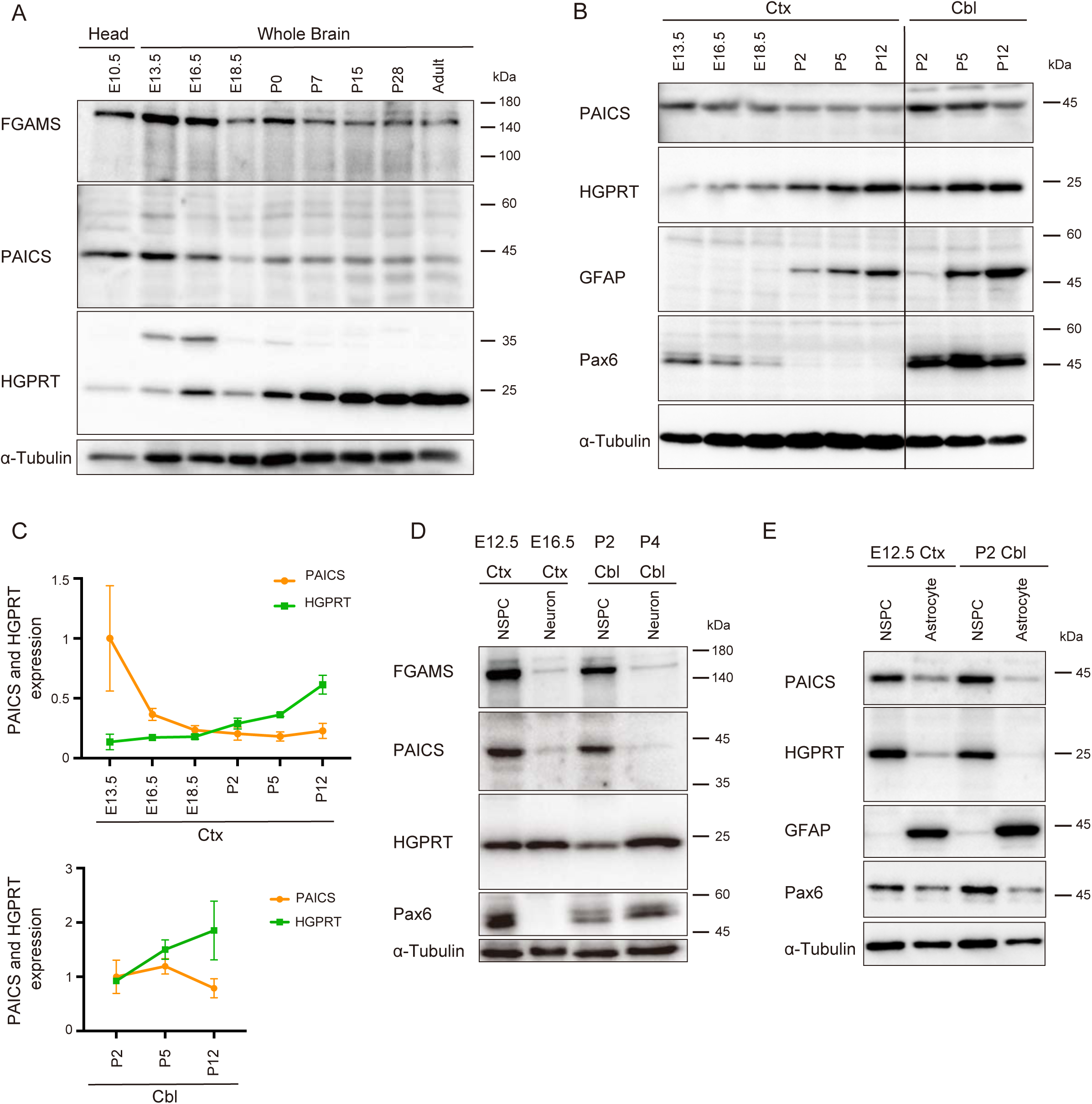
Developmental expression of the enzymes of the *de novo* and salvage purine synthesis pathways in the nervous system. PAICS, FGAMS, and HGPRT expression was assessed via western blotting using embryonic and postnatal brains (A–C) or primary cultured cells (D–E). (A) Developmental changes in the expression of each enzyme in whole-brain lysates prepared from different developmental stages (E10.5–adult). (B) PAICS, HGPRT, GFAP, and Pax6 expression in the cerebral cortex (ctx) (E13.5–P12) or cerebellum (cbl) (P2–P12). (C) The line charts representing the expression of PAICS and HGPRT during ctx (upper) and cbl development (bottom). The bands obtained in three independent mouse brains in each developmental stage and the brain region of blots were quantified using α-Tubulin as an internal standard, and the mean values were plotted for each age. (D) Expression of FGAMS, PAICS, HGPRT, and Pax6 in cultured NSPCs or differentiated neurons prepared from the ctx (E12.5 or E16.5) and cbl (P2 or P4). (E) Expression of PAICS, HGPRT, GFAP, and Pax6 in primary cultured NSPCs or astrocytes. α-Tubulin was used as a loading control (bottom panels).

We next analyzed the expression of these enzymes in primary cultured NSPCs isolated from E12.5 cerebral cortices and the external granular layer (EGL) of the P2 cerebellum. Figure 2D shows that PAICS and FGAMS were abundantly expressed in Pax6^+^ NSPCs derived from these two brain regions. In contrast, PAICS and FGAMS expression was decreased in primary cultured neurons (Fig. 2D) isolated from E16.5 cerebral cortices or P4 cerebella and cultured for 10 div until fully differentiated. These observations appear consistent with the *in vivo* expression profile of FGAMS and PAICS along brain development (Fig. 2A–C). On the other hand, HGPRT was expressed in cultured NSPCs as well as the differentiated neurons (Fig. 2D). We further analyzed the expression of these enzymes in primary cultured GFAP^+^ astrocytes, which were differentiated from the embryonic cerebral cortex and postnatal cerebellum. However, the expression of both HGPRT and PAICS was barely detectable in astrocytes (Fig. 2E). The purity of the NSPCs and astrocytes used in these assays was confirmed by the expression of Pax6, a marker for embryonic apical progenitor cells/radial glial cells (RGCs), and GFAP, a marker for astrocyte, respectively (Fig. 2D, E). In the cerebellum, the Pax6 protein is expressed in differentiated granular cells as well as cerebellar NSPCs (Yamasaki et al., 2001). Consistently, we detected Pax6 in the cultured cerebellar neurons (Fig. 2D). Together, these results suggested that the *de novo* pathway is preferentially driven during the embryonic stage when NSPC proliferation occurs, and that purine metabolism switches from the *de novo* pathway to the salvage pathway over brain development.

### Expression of purine synthesis enzymes in the developing and adult CNS

The distribution and localization of PAICS, FGAMS, and HGPRT were examined immunohistochemically in the developing and matured mouse brain. At E13.5, three enzymes were strongly expressed in the VZ/SVZ of the cerebral cortex, where NSPCs localize (Fig. 3A–C). These immunoreactivities in the VZ/SVZ considerably decreased until E18.5 and P0 (Fig. 3A–F, J–L), decreasing until disappearing in the SVZ of the adult cerebral cortex (Fig. 3P–R). The development of the cerebellum proceeds rapidly from late embryonic stages. Granule neurons are produced from NSPCs in the superficial EGL covering the cerebellar cortex and migrate to form the internal granule layer (IGL) (Yacubova & Komuro, 2003). We observed robust HGPRT immunoreactivity in the EGL at E18.5 and P0; meanwhile, the PAICS and FGAMS expression level in the EGL was relatively weak (Fig. 3G–I, M–O, arrows). These observations might indicate that embryonic NSPCs actively drive purine synthesis pathways to meet the high demand at the beginning; as neurogenesis progresses though, the activity of the purine synthesis pathway would decline. We further detected HGPRT expression in CD31^+^ endothelial cells of blood vessels in the brain parenchyma from E13.5 to P0.5 (Fig. 3S, Extended Data Fig. 3-1), but HGPRT was not observed in blood vessels in the adult brain. These results suggested a specific role of the salvage pathway in vasculogenesis in the embryonic brain.

**Figure 3.**
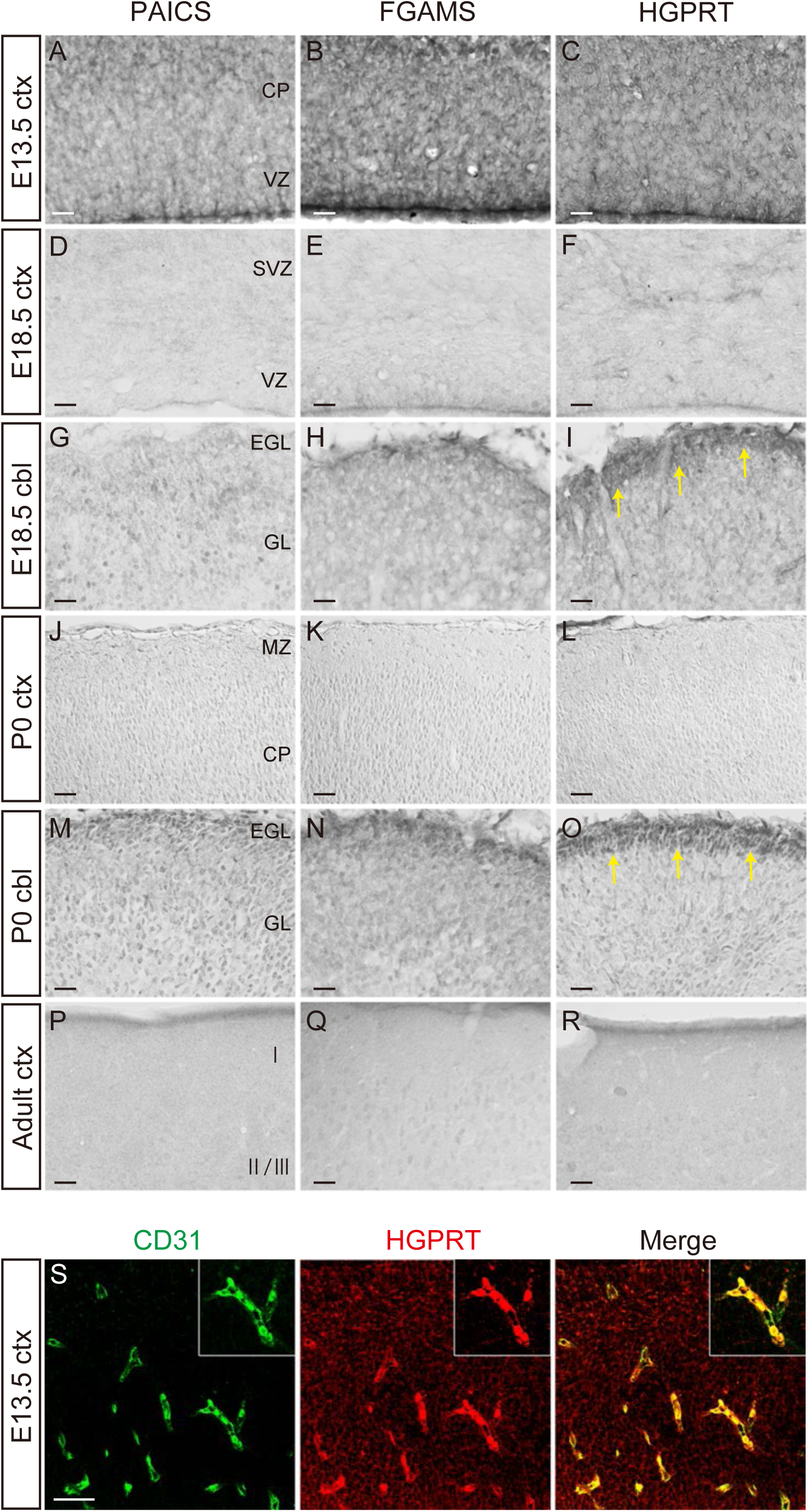
Expression profile of purine synthesis enzymes in the developing cerebral cortex and cerebellum. (A–R) Immunostaining with anti-PAICS (A, D, G, J, M, P), anti-FGAMS (B, E, H, K, N, Q), or anti-HGPRT (C, F, I, L, O, R) antibodies in developmental ctx and cbl. Frozen sections of E13.5 ctx (A–C), E18.5 ctx (D–F), E18.5 cbl (G–I), P0 ctx (J–L), P0 cbl (M–O), and adult ctx (P–R) showing the distribution of PAICS, FGAMS, HGPRT-positive cells. Arrows indicate the HGPRT expression in EGL. (S) Representative confocal image of the E13.5 cerebral cortex double immunostained with the anti-CD31 (*green*) and anti-HGPRT (*red*) antibodies. The inset shows a higher magnification of CD31^+^ HGPRT^+^ endothelial cells. CP, cortical plate; VZ, ventricular zone; SVZ, subventricular zone; EGL, external granule cell layer; GL, internal granule cell layer; MZ, marginal zone; I, cortical layer I; Ⅱ/Ⅲ, cortical layers Ⅱ and Ⅲ. Scale bars, 100 µm in (A‒R); 50 µm in (S).

In the adult brain, PAICS, FGAMS, and HGPRT were widely detected in neuronal somata and neuropil of discrete areas (Extended Data Fig. 3-2 and 3-3). In the brain stem, the locus coeruleus, Edinger-Westphal nucleus, and vestibular nucleus expressed these three enzymes. Moreover, strong immunoreactivities for PAICS and FGAMS were differentially observed in several nuclei of the brain stem, including the lateral cerebellar, red, facial, medial vestibular, and ambiguous nucleus (Extended Data Fig. 3-2 and 3-3). On the other hand, HGPRT was preferentially expressed in numerous nuclei in the forebrain and diencephalon, including the posterior interlaminar thalamic nucleus, polymorph layer of the dentate gyrus, arcuate nucleus, dorsomedial hypothalamic nucleus, zona incerta, lateral septal nuclei, and neurons in the nigrostriatal bundle (Extended Data Fig. 3-2 and 3-3). These findings implied that the purine pathway to be activated depends on the developmental stage and brain region involved.

### Inhibiting *de novo* purine synthesis affects NSPC proliferation

We sought to investigate the effects on brain development of purine synthesis inhibition during a specific period in embryonic and postnatal life. To this end, using genetically engineered animals, such as knockout mice and conditional knockout mice of purine synthesis enzymes, was limited in their ability to transiently suppress the enzyme activity in the mid or late embryonic period, times of active neurogenesis in the cerebral cortex. In this study, we used several inhibitors that target specific enzymes of purine synthesis pathways. These inhibitors allowed us to target a specific embryonic period and to easily adjust the degree of inhibition during corticogenesis depending on inhibitor concentration.

First, the role of purine pathways in NSPCs was explored using various specific inhibitors for the *de novo* and salvage enzymes. Accordingly, primary cultured NSPCs prepared from E12.5 cerebral cortex were incubated for 48 h with or without the following inhibitors (Fig. 1): MMF, an inhibitor of the inosine monophosphate dehydrogenase, which is the rate-limiting enzyme in *de novo* synthesis of guanosine nucleotides (Allison, 2005) or forodesine hydrochloride, an inhibitor of the purine nucleoside phosphorylase mediating the salvage pathway (Evans et al., 2018). Immunostaining with anti-Nestin, an NSPC marker and anti-BrdU revealed that MMF treatment significantly reduced the number of Nestin^+^ BrdU^+^ NSPCs, while forodesine treatment did not affect NSPC proliferation (Fig. 4A–D). MMF treatment led to no morphological changes or enhanced differentiation into neurons in Nestin^+^ NSPCs, suggesting that MMF exclusively affects the mitotic potential of NSPCs. Similar results were obtained in the NSPCs isolated from the cerebellar EGL (Extended Data Fig. 4-1). Next, we confirmed that the reduced NSPC proliferation was caused by repression of the *de novo* pathway using PAICS shRNAs. The silencing effect and specificity of PAICS shRNAs (#1 and #3) were validated in our previous study (Yamada et al., 2020). Consistent with the inhibitory effect of MMF on NSPC division, knocking down *PAICS* significantly decreased the number of BrdU-incorporated NSPCs (Paics shRNA #1, 37%, n = 100; Paics shRNA #3, 6%, n = 100, arrows) relative to the control (control shRNA, 67%, n = 100, arrowhead) (Fig. 4E–G). Based on these results, we concluded that activation of the *de novo* pathway is critical for NSPC proliferation.

**Figure 4.**
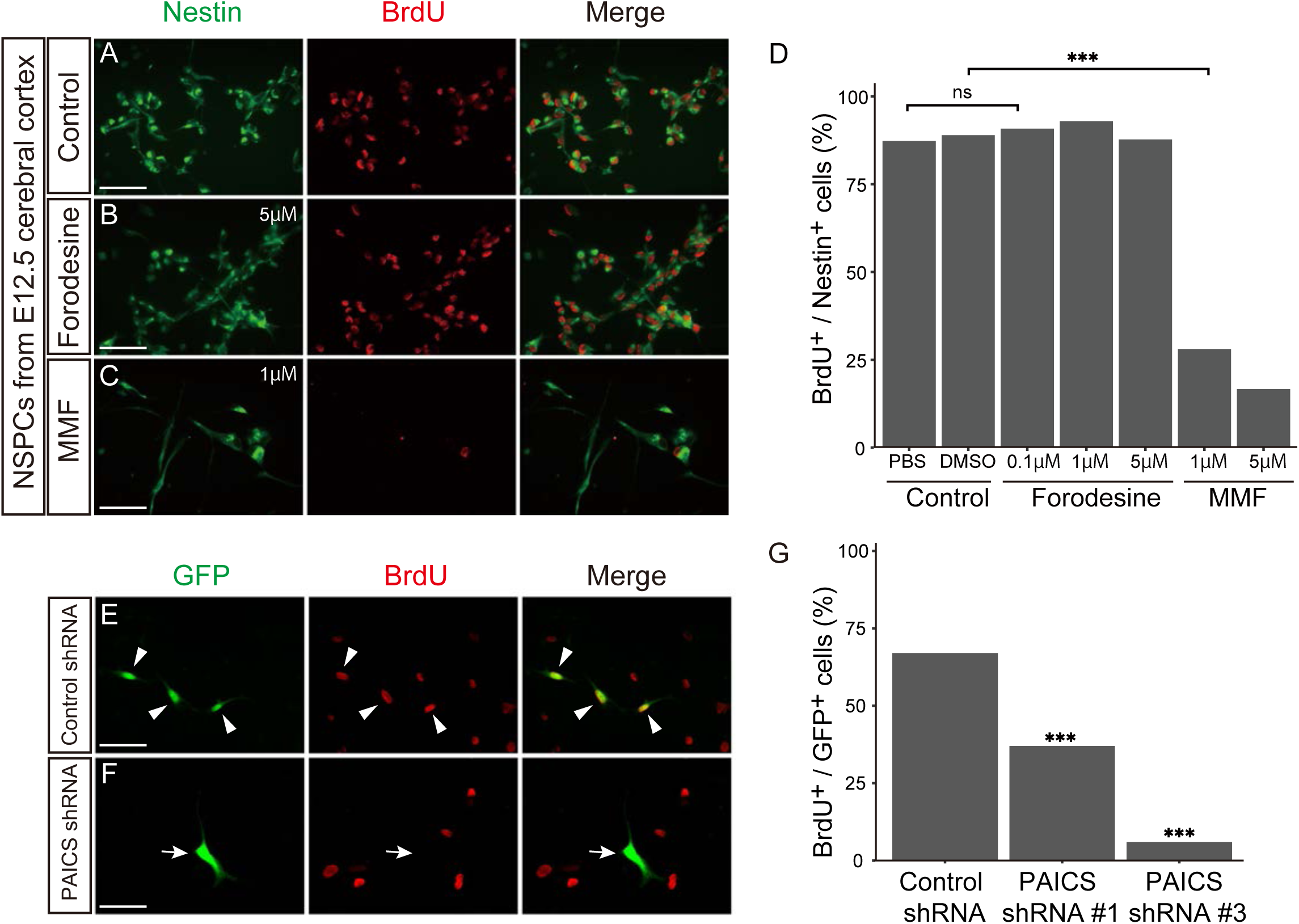
Inhibition of *de novo* purine pathway disturbs NSPC proliferation *in vitro*. (A–D) Primary cultured NSPCs from the E12.5 telencephalon were treated for 48 h with PBS or DMSO as control (A), forodesine (0.1, 1, 5 µM) (B), or mycophenolate mofetil (MMF) (1, 5 µM) (C), followed by BrdU labeling for 24 h. NSPCs were immunostained with anti-Nestin (*green*) and anti-BrdU (*red*) antibodies. (D) Quantified comparison of the number of the BrdU^+^ Nestin^+^ cells to total Nestin^+^ cells. ns, not significant; ***p < 0.001, chi-square tests with Holm–Bonferroni correction. control PBS, n = 326; control DMSO, n = 346; forodesine 0.1 µM, n = 435; forodesine 1 µM, n = 283; forodesine 5 µM, n = 310; MMF 1 µM, n = 114; MMF 5 µM, n = 72. (E–G) Primary cultured NSPCs were electroporated with control shRNA (E) or PAICS shRNAs (#1 or #3) (F) together with EGFP-expression plasmid (*green*), followed by labeling with BrdU (*red*) for 24 h. Arrowheads indicate GFP^+^ BrdU^+^ proliferating NSPCs. The arrow indicates EGFP^+^ but BrdU^−^ cells. (G) The numbers of GFP^+^ BrdU^+^ cells were compared to the total GFP^+^ cells using the chi-square tests with Holm– Bonferroni correction. ns, not significant; ***p < 0.001. control shRNA, n = 100; PAICS shRNA#1, n = 100; PAICS shRNA#3, n = 100. Scale bars, 50 µm.

#### The development of the cerebellum requires both purine synthesis pathways

Next, we examined the significance of the purine pathways on *in vivo* neurogenesis in the cerebellum. To this end, embryos at E16.5 or pups at P2 mice were treated with forodesine, MMF, or MTX. MTX is known to impede the *de novo* purine pathway at multiple steps by inhibiting the dihydrofolate reductase, which catalyzes the folate biosynthesis pathway (Fig. 1) (Cronstein, 1997). Gross anatomical analysis two or four days after inhibitor administration showed no significant brain dysplasia. However, immunostaining with BrdU and Ki67, which marks cells with proliferative capacity, showed that the fraction of BrdU^+^ and Ki67^−^ cells increased in the E18.5 cerebellar IGL treated with forodesine, MMF, and MTX compared with control (Fig. 5A–E) (control, 20.6 ± 2.7%, n = 7; forodesine, 40.3 ± 2.7%, n = 8; MMF, 34.4 ± 2.9%, n = 7; MTX, 49.6 ± 3.2%, n = 9). Comparable results were observed in the P6 cerebellar IGL treated with forodesine or MMF (Fig. 5F–I) (control, 27.1 ± 1.5%, n = 9; forodesine, 38.4 ± 3.2%, n = 7; MMF, 42.8 ± 2.7%, n = 9). These findings suggest that inhibition of purine synthesis in the cerebellum may accelerate the exit from the cell cycle in NSPCs, affecting cell fate determination and leading to impaired cerebellar development. Considering the abundant HGPRT expression in EGL (Fig. 3I, O), proper development of the cerebellum requires the salvage as well as the *de novo* pathways.

**Figure 5.**
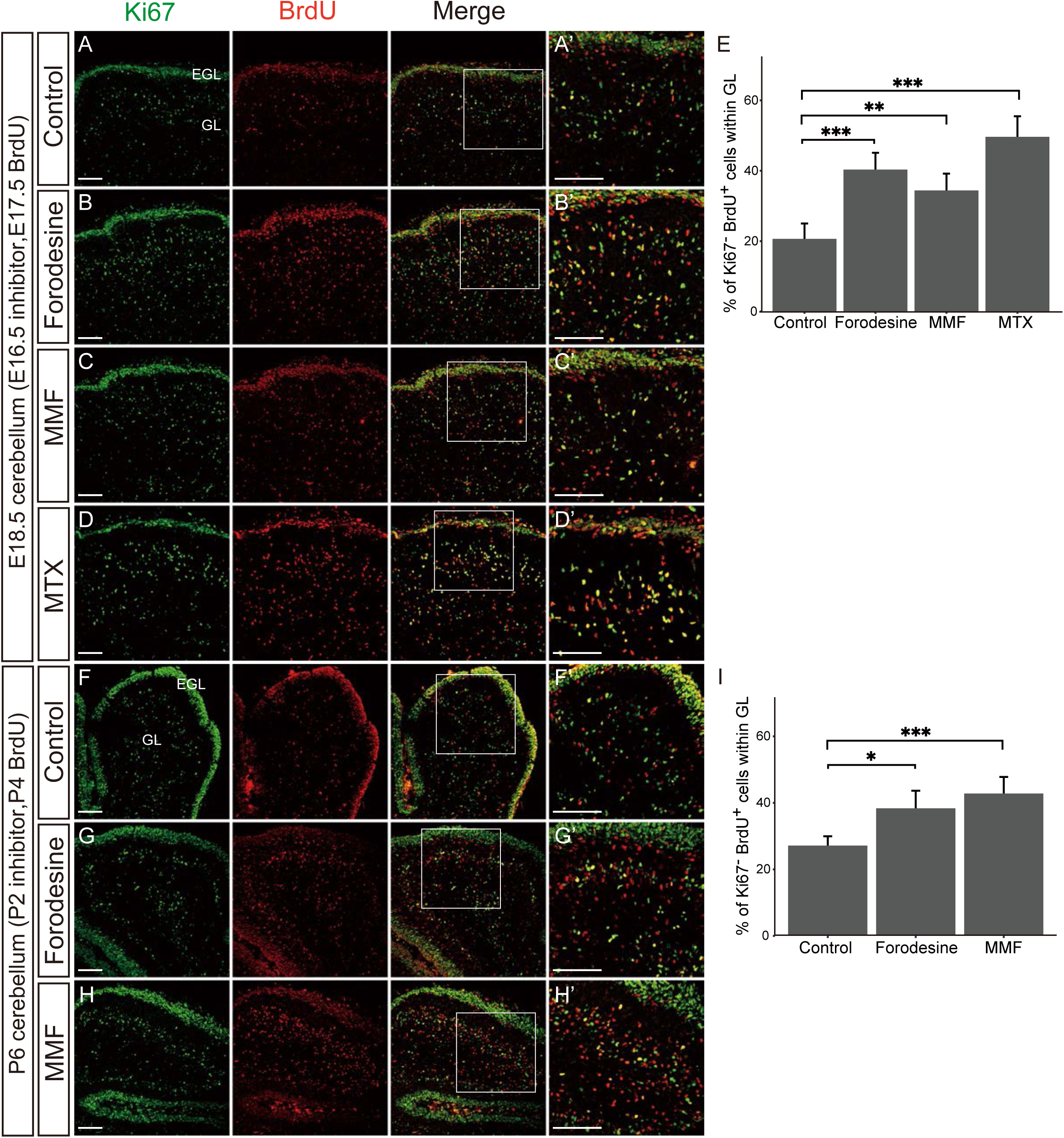
The development of the cerebellum is cooperatively regulated by both purine synthesis pathways. (A–D) Confocal images of the E18.5 cerebellum stained with the Ki67 (*green*) and BrdU (*red*). Control DMSO (A), forodesine (B), MMF (C), or methotrexate (MTX) (D) was administered at E16.5, followed by labeling with BrdU at E17.5, and the cerebellum was analyzed at E18.5. (A’–D’) Higher magnification of A–D. (E) Quantified comparison of the percentage of Ki67^−^ BrdU+ cells to total BrdU^+^ cells in GL at E18.5. (F–H) Confocal images of the P6 cerebellum stained with the Ki67 (*green*) and BrdU (*red*). Control DMSO (F), forodesine (G), or MMF (H) was administered to P2 pups, followed by BrdU labeling at P4, and the cerebellum was immunostained at P6. (F’–H’) Higher magnification of F–H. (I) Quantified comparison of the percentage of Ki67^−^ BrdU^+^ cells to the total BrdU^+^ cells in the GL at P6. Data are presented as means ± SEM. *p < 0.05, **p < 0.01, ***p < 0.001, Welch’s *t*-test followed by Holm–Bonferroni correction. EGL, external granule cell layer; GL, internal granule cell layer. Scale bars, 100 µm.

### MMF suppresses NSPC proliferation and delays neuron migration in the cerebral cortex

To assess the impact of the purine pathway on neurogenesis in the embryonic cerebral cortex, MMF or forodesine was administered at E12.5, followed by BrdU administration at E13.5, and embryo analysis at E14.5. In the control, many BrdU-incorporated NSPCs exited the cell cycle and left the SVZ/VZ; about half of BrdU^+^ cells were dispersed within the intermediate zone (IZ) and cortical plate (CP) at E14.5 (Fig. 6A, D) (CP, 8.2 ± 1.1%; IZ, 42.9 ± 2.0%; SVZ/VZ, 48.9 ± 1.3%, n = 9). In contrast, MMF-treated embryos exhibited a substantial accumulation of BrdU^+^ cells in the SVZ/VZ, with few BrdU^+^ cells being located in the IZ/CP (Fig. 6C, D) (CP, 0.1 ± 0.1%; IZ, 27.7 ± 3.5%; SVZ/VZ, 72.2 ± 3.5%, n = 9), indicating suppression of radial migration of newborn neurons. On the other hand, forodesine treatment had only a small effect on neuron migration (Fig. 6B, D) (CP, 2.5 ± 0.8%; IZ, 40.4 ± 3.1%; SVZ/VZ, 57.1 ± 3.1%, n = 9). To determine whether the BrdU^+^ cells remaining in the VZ/SVZ had lost their mitotic potential, we performed immunostaining with an antibody to phospho-histone H3 (pH3), which marks cells in late G2-M phase. Compared to controls, the number of pH3^+^ NSPCs in the VZ facing the lateral ventricles was significantly reduced in MMF-treated embryos (Fig. 6E, G, H, arrows) (control, 31.1 ± 3.5, n = 11; MMF, 19.0 ± 1.4%, n = 9). Meanwhile, the number of pH3^+^ NSPCs in the VZ remained unchanged in forodesine-treated embryos (Fig. 6F, H) (34.2 ± 1.9%, n = 9). Immunostaining with anti-PAICS, FGAMS, and HGPRT showed that the expression and subcellular localization of each enzyme were not affected by treatment with these inhibitors, as all these enzymes were expressed in the cytoplasm in the VZ/SVZ (Fig. 6I–Q). Furthermore, treatment with these inhibitors did not induce early premature differentiation into astrocytes, as demonstrated by immunostaining with anti-GFAP (Extended Data Fig. 6-1). These results suggested that inhibition of the *de novo* pathway, but not of the salvage pathway, suppressed NSPC mitosis and the subsequent neuron migration from the VZ.

**Figure 6.**
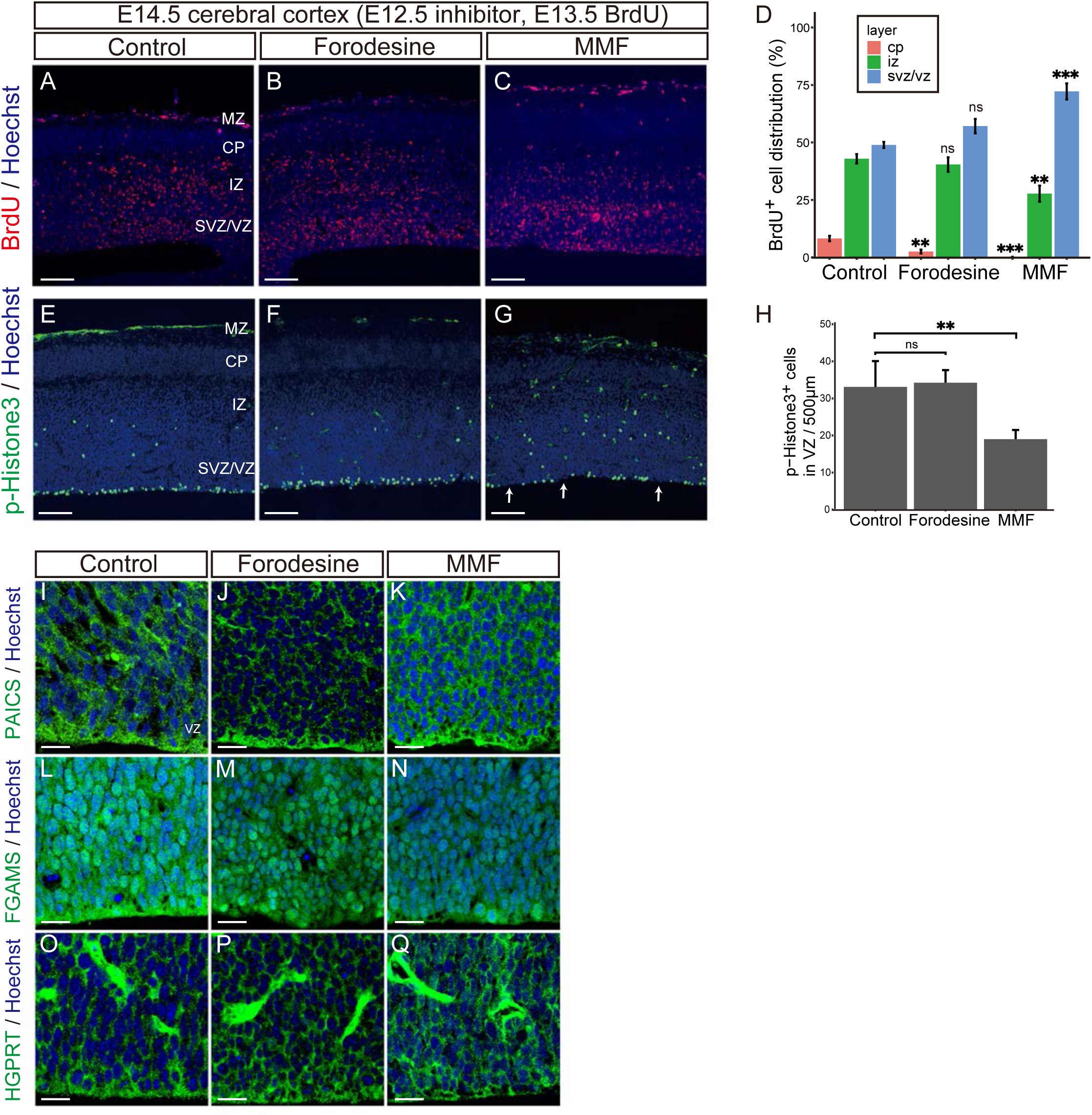
Formation of the cerebral cortex mainly depends on the *de novo* purine pathway. (A–H) E12.5 embryos were treated with control DMSO (A, E), forodesine (B, F), or MMF (C, G), followed by BrdU labeling, and analyzed at E14.5. Horizontal frozen sections were immunostained with anti-BrdU antibody (*red*) (A–C) or phospho-histone H3 (pH3) antibody (*red*) (E–H). Arrows in (G) indicate pH3^−^ VZ areas. (D) Quantification of the percentage of BrdU^+^ cells distribution in each layer (CP, IZ, SVZ/VZ). (H) Number of pH3^+^ cells/500 µm width of the VZ region. Data are presented as means ± SEM. ns, not significant, **p < 0.01, ***p < 0.001, Welch’s *t*-test followed by Holm–Bonferroni correction. (I–Q) Immunostaining of PAICS (I–K), FGAMS (L–N), and HGPRT (O–Q) in the VZ/SVZ area of E14.5 embryos treated with DMSO (I, L, O), forodesine (J, M, P), or MMF (K, N, Q). Nuclei were stained with Hoechst dye (*blue*). MZ, marginal zone; CP, cortical plate; IZ, intermediate zone; SVZ/VZ, subventricular zone/ventricular zone. Scale bars, 100 µm in (A–C) and (E–G); 20 µm in (I–Q).

### *De novo* purine pathway controls mTORC1/S6K/S6 signaling

We attempted to elucidate the molecular mechanism by which the *de novo* purine synthesis pathway regulates cortical development. Among various intracellular signaling pathways, we focused on mTOR, a serine/threonine kinase involved in the proliferation and growth of cells through the regulation of numerous cellular processes (Szwed et al., 2021). Previous studies have shown that mTORC1 stimulates purine nucleotide production (Ben-Sahra et al., 2016) as well as the inhibitory effect of purine nucleotide depletion on mTORC1 signaling (Hoxhaj et al., 2017). In addition, mTORC1 mutation affects the development of the cerebrum, suggesting a role of mTORC1 signaling in NSPCs (Tarkowski et al., 2019; Andrews et al., 2020). mTORC1 signaling includes two downstream cascades, namely eukaryotic initiation factor 4E (eIF4E)/binding protein 1 (4E-BP1) and S6K/S6 (Fig. 7C) (Ben-Sahra et al., 2016; Morita et al., 2013). These two downstream cascades play an essential role in protein translation. Specifically, mTORC1 pS6K, which further phosphorylates various substrate proteins, including ribosomal subunit S6, thereby promoting translation. However, the molecular interplay between the two mTORC1 signaling cascades and purine metabolism in cortical development remains unclear.

**Figure 7.**
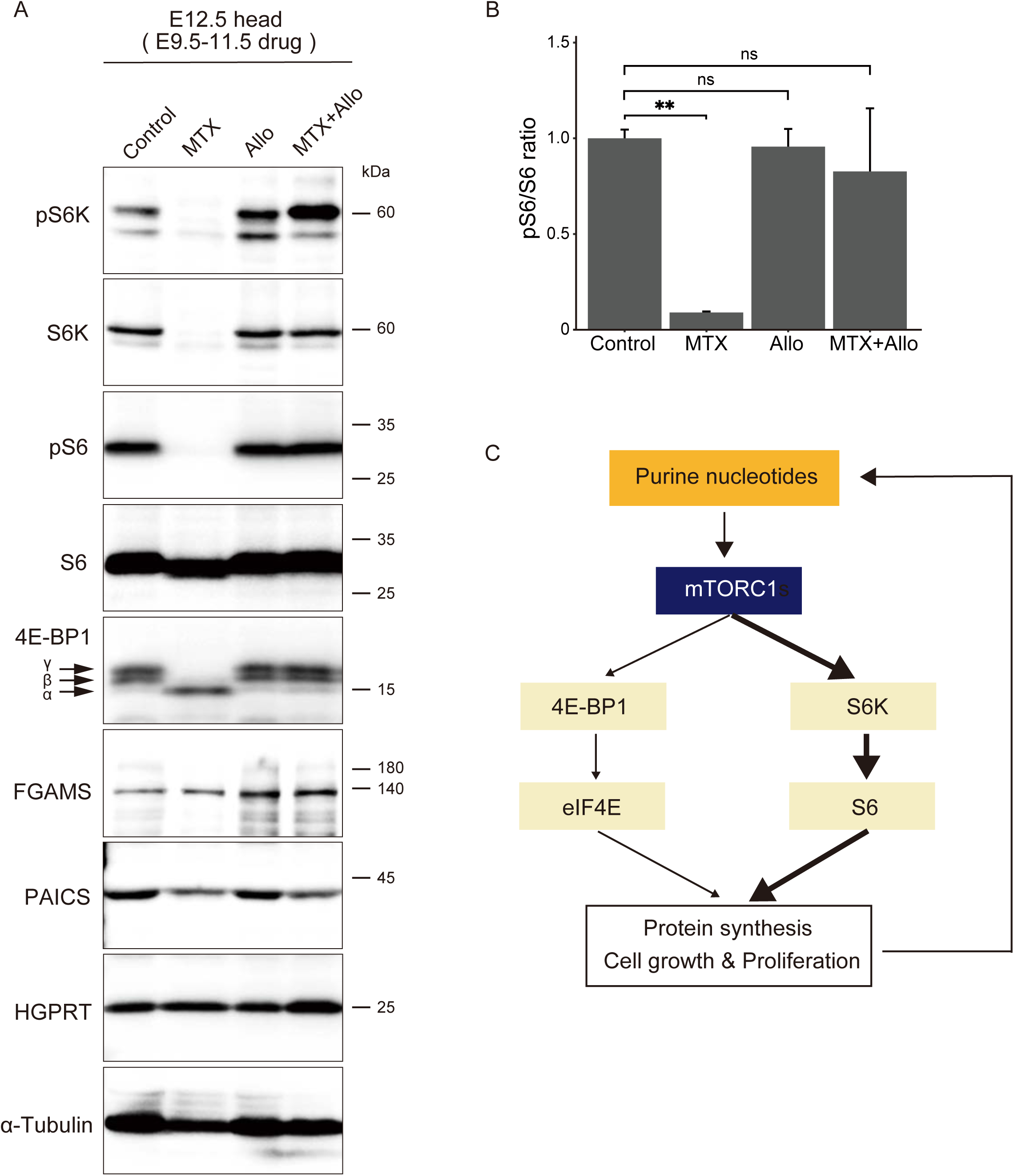
The *de novo* purine pathway affects mTORC1/S6K/S6 signaling. (A) Immunoblot analysis of E12.5 brains treated with each inhibitor at E9.5–E11.5. Each panel presents the expression of mTOR signaling proteins (pS6K, S6K, pS6, S6, and 4E-BP1) and purine synthesis enzymes (PAICS, FGAMS, and HGPRT). Arrows denote the α, β, and γ isoforms of 4E-BP1. The blot was reprobed with α-Tubulin antibody (bottom) to examine quantitative protein loading. (B) Quantified comparison of the pS6/S6 ratio. Data are presented as the mean ± SEM. ns, not significant, **p < 0.01, Welch’s *t*-test followed by the Holm–Bonferroni correction. (C) Schematic diagram of the relationship between mTOR signaling and purine nucleotides.

Brain lysates were prepared from E12.5 embryos continuously treated with MTX, allopurinol, or MTX plus allopurinol, and the expression of the downstream proteins of mTORC1 signaling was examined. Allopurinol is an inhibitor of xanthine oxidase, which converts xanthine to uric acid, thus promoting purine synthesis via the salvage pathway (Fig. 1) (Day et al., 2017). PAICS expression was decreased by MTX, but HGPRT expression showed no change; thus, MTX specifically inhibits *de novo* purine synthesis. For the S6K/S6 signaling cascade, we analyzed the expression of S6K, pS6K, S6, and pS6. As shown in Figure 7A, pS6K, S6K, and pS6 expression was completely inhibited by MTX treatment. Compared with the control, S6 protein expression remained unchanged in MTX-treated brains, indicating that MTX completely inhibits S6 phosphorylation and activation in the embryonic brain (Fig. 7B). These severe MTX-induced impairments of the S6K/S6 signaling cascade were restored by the concurrent administration of allopurinol and MTX (Fig. 7A), which eventually restored the phosphorylation level of S6 to control levels (Fig. 7B). Regarding another downstream pathway of mTORC1, 4E-BP1 inhibits cap-dependent translation by binding to the translation initiation complex eIF4E (Sun et al., 2019). Three 4E-BP1 isoforms (α, β, and γ) exist in mammalian cells that represent the phosphorylation status of 4E-BP1, with α being hypophosphorylated and β/γ being hyperphosphorylated (Gingras et al., 1998). We found a mobility shift in MTX-treated brains from 4E-BP1-β/γ to 4E-BP1-α (arrows in Fig. 7A, Extended Data Fig. 7-1B), although quantitative analysis revealed little difference in total 4E-BP1 protein expression (Extended Data Fig. 7-1A). A previous report consistently indicated that the accumulation of 4E-BP1-α was induced by MTX treatment in cultured HeLa cells (Hoxhaj et al., 2017). Considering that 4E-BP1-α can efficiently interact with and inhibit eIF4E (Gingras et al., 1998), the MTX-induced dephosphorylation of 4E-BP1 might repress eIF4E-dependent translation in the embryonic brain.

These results indicate that the *de novo* purine synthesis pathway is strongly associated with mTORC1/S6K/S6 signaling and is related to mTORC1/4E-BP1/eIF4E signaling to some extent. As the activation of the salvage pathway by allopurinol reversed the inhibition of mTORC1/S6K/S6 signaling, we considered that a steady-state level of purines supplied by the *de novo* pathway is essential for the maintenance of mTORC1 signaling within NSPCs *in vivo* (Fig. 7C).

### Activation of mTOR signaling abrogates inhibition of the *de novo* pathway

Inhibition of *de novo* synthesis in NSPCs often induces cell death in addition to reducing mitotic potential (Fig. 4). We determined whether this apoptosis could be rescued by mTOR activation. For this purpose, NSPCs were treated with purine inhibitors along with MHY1485, an activator of the mTOR signaling pathway (Choi et al., 2012). The primary cultured NSPCs prepared from the E12.5 cerebral cortex were co-incubated with or without MHY1485, and the apoptotic rate was evaluated via immunostaining for cleaved-caspase3 at 6 or 48 h. Treatment with MMF resulted in a significant induction of apoptosis in Nestin^+^ NSPCs, whereas treatment with forodesine showed no effect (Fig. 8A, B, C, E). This deleterious effect of MMF was entirely abolished by co-treatment with MHY1485 (Fig. 8C, E). Notably, neither MMF nor forodesine affected cell death in GFAP^+^ astrocytes (Fig. 8F–J). These results suggested that apoptosis caused by inhibition of the *de novo* pathway does not simply result from impaired proliferative ability; instead, it represents a phenomenon attributable to cellular characteristics unique to NSPCs, such as multipotency, wherein the activation of mTORC1 signaling is critical for maintenance or survival.

**Figure 8.**
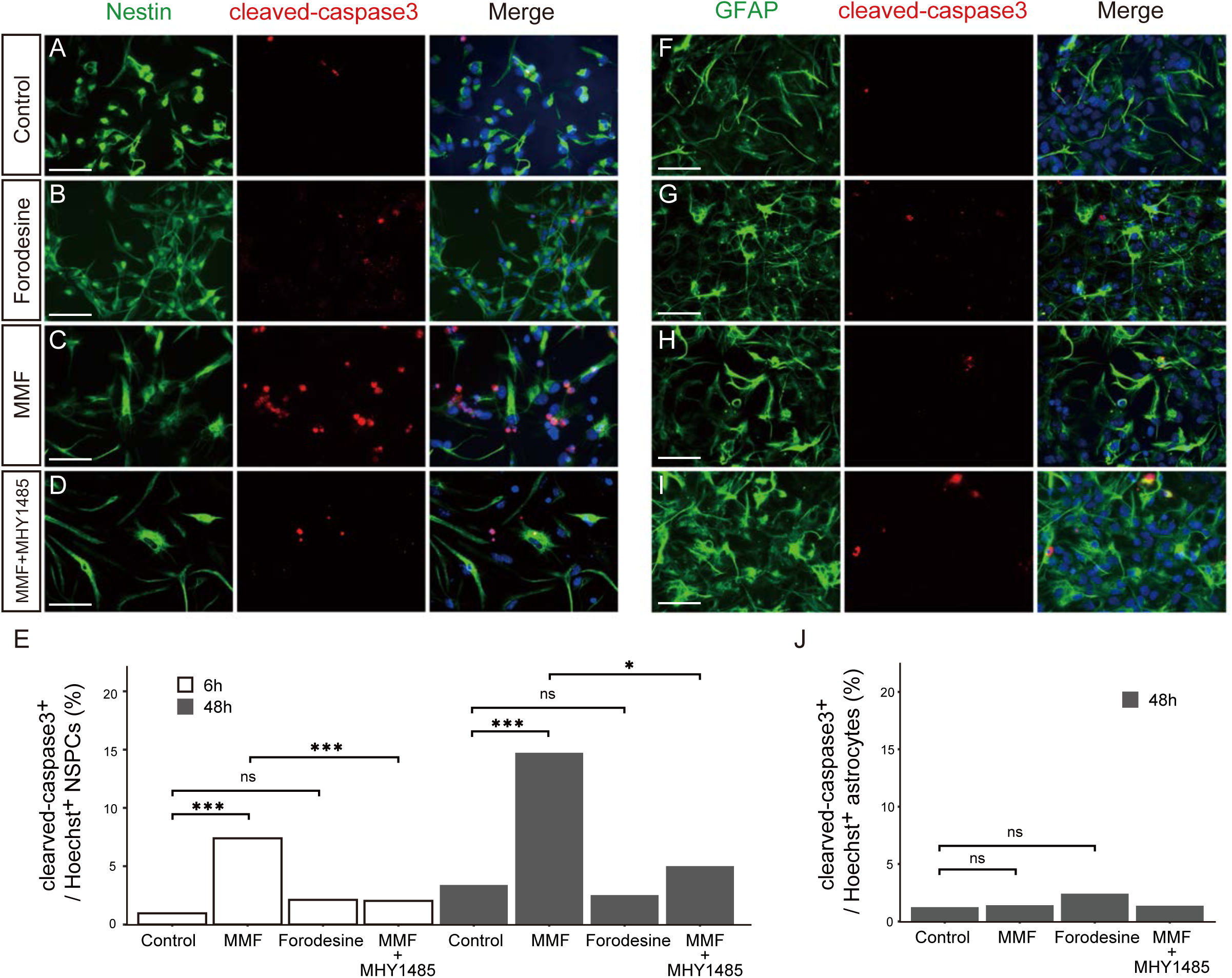
Inhibition of the *de novo* purine synthetic pathway causes apoptosis in NSPCs but not in astrocytes. (A–E) Primary cultured NSPCs from the E12.5 telencephalon were treated for 48 h with DMSO as a control (A), forodesine (50 µM) (B), MMF (10 µM) (C), or MMF and MHY1485 (10 µM) (D). NSPCs were immunostained with anti-Nestin (*green*) and anti-cleaved-caspase3 (*red*) antibodies. (E) Quantification of apoptotic NSPCs. The number of cleaved-caspase3^+^ Nestin^+^ cells was counted and represented as a ratio to the number of Hoechst^+^ Nestin^+^ cells at 6 (white bar) or 48 h (black bar). (F–J) Astrocytes differentiated from NSPCs were treated for 48 h with DMSO (F), forodesine (50 µM) (G), MMF (10 µM) (H), or MMF and MHY1485 (10 µM) (I). Astrocytes were immunostained with anti-GFAP (*green*) and anti-cleaved-caspase3 (*red*) antibodies. (J) Quantification of apoptotic astrocytes. The number of cleaved-caspase3^+^ GFAP^+^ cells was counted and represented as a ratio to the number of Hoechst^+^ GFAP^+^ cells. ns, not significant, *p < 0.05, ***p < 0.001, chi-squared test with the Holm–Bonferroni correction. control, n = 313 (E, 48 h), 236 (E, 6 h), 322 (J); forodesine, n = 159 (E, 6 h), 237 (E, 48 h), 746 (J); MMF, n = 284 (E, 6 h), 224 (E, 48 h), 354 (J); MMF and MHY1485, n = 100 (E, 6 h), 148 (E, 48 h), 292 (J). Scale bars, 50 µm.

### Inhibiting *de novo* pathway causes forebrain-specific malformations

To determine how the chronic inhibition of the purine pathway impacts brain formation *in vivo*, different inhibitors (MMF, forodesine, MTX, and allopurinol) were continuously administered to embryos from E9.5 to E11.5, with or without MHY1485, and their brain architecture was analyzed at E12.5. As shown in Figure 9, embryos treated with forodesine, MHY1485, or allopurinol alone appeared to follow normal brain development (Fig. 9A, C, E, Extended Data Fig. 9-1). On the other hand, a drastic anomaly was reproducibly observed in MMF-treated brains, in which the lateral ventricles widely opened into the third ventricle, probably due to forebrain hypoplasia (Fig. 9B). The combined administration of MMF and MHY1485, an activator of the mTOR signal, could restore this gross malformation (Fig. 9D). MTX-treated embryos exhibited a more severe defect with complete loss of the forebrain region (Fig. 9G). This systematic abnormality was extremely severe and could not be reversed by the combined administration of MTX and allopurinol, thereby enhancing the salvage pathway (Fig. 9H). These observations further supported that the *de novo* purine synthesis pathway is predominantly involved in the development of the cerebral cortex, as the *de novo* synthesis inhibition impacted forebrain formation in early neural development.

**Figure 9.**
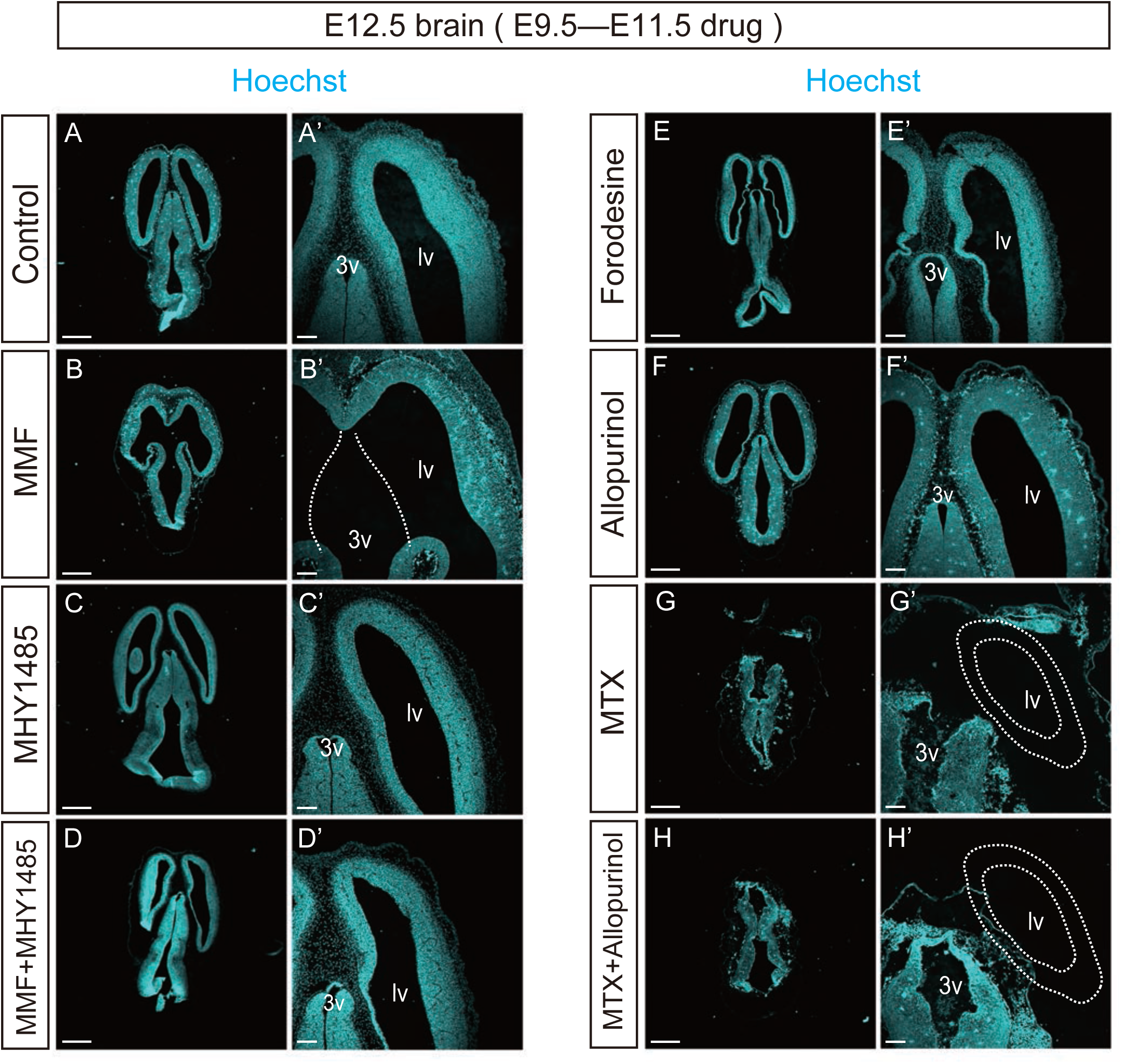
Embryonic brain malformations induced by inhibiting *de novo* purine pathway. Horizontal frozen sections of E12.5 brains. Control DMSO (A), MMF (B), MHY1485 (C), MMF and MHY1485 (D), forodesine (E), allopurinol (F), MTX (G), or MTX and allopurinol (H) were successively administered during E9.5–E11.5 and brains were harvested at E12.5. Sections were stained with Hoechst dye (cyan). (A’–F’) Magnified views of the forebrain of A–H. lv, lateral ventricles; 3v, 3rd ventricle. Scale bars, 500 µm in (A–H), 100 µm in (A’–H’).

We performed a detailed histological analysis to gain insights into the cellular mechanism of the brain abnormality caused by the suppression of *de novo* purine synthesis. As shown in Figure 10, MMF-treated embryos showed a significant increase in the number of cleaved-caspase3^+^ apoptotic cells in the rostral region of the cerebral cortex, whereas a considerably lower number of apoptotic cells were detected in the caudal forebrain region (Fig. 10A, B, AF) (control, 7.2 ± 1.2, n = 4; MMF rostral area, 47.3 ± 7.7, n = 8; MMF caudal area, 18.8 ± 2.7, n = 8). Co-treatment with MHY1485 successfully rescued the morphological phenotype induced by MMF and reduced the number of apoptotic cells, thereby restoring the normal histological structure of the brain (10.8 ± 2.9, n = 9, Fig. 10C, AF). As illustrated in Figure 9G, the consecutive administration of MTX led to severe cerebral malformations accompanied with an increase in cell death and positivity for cleaved-caspase3. This cell death was entirely ameliorated by treatment with MHY1485, as determined by western blot using anti– cleaved-caspase3. (Fig. 10AH, AI), implicating that the MTX-induced brain phenotype depends on the mTORC1 pathway. In contrast, neither the continuous inhibition of the salvage pathway by forodesine nor the activation of the salvage pathway by allopurinol had any effect on brain formation (Fig.10D, E). In addition, forodesine treatment did not affect cell death (Fig.10D, AF) (16.4 ± 2.2, n = 6). Interestingly, allopurinol treatment resulted in a marked increase in the number of apoptotic caspase3^+^ cells throughout the cerebral cortex (Fig. 10E, AF) (42.9 ± 5.6, n = 9). Because uric acid is a major antioxidant and its neuroprotective effects reduce the risk of neurological diseases (Kachroo & Schwarzschild, 2014), suppressing uric acid production with allopurinol can potentially affect the developing brain.

**Figure 10.**
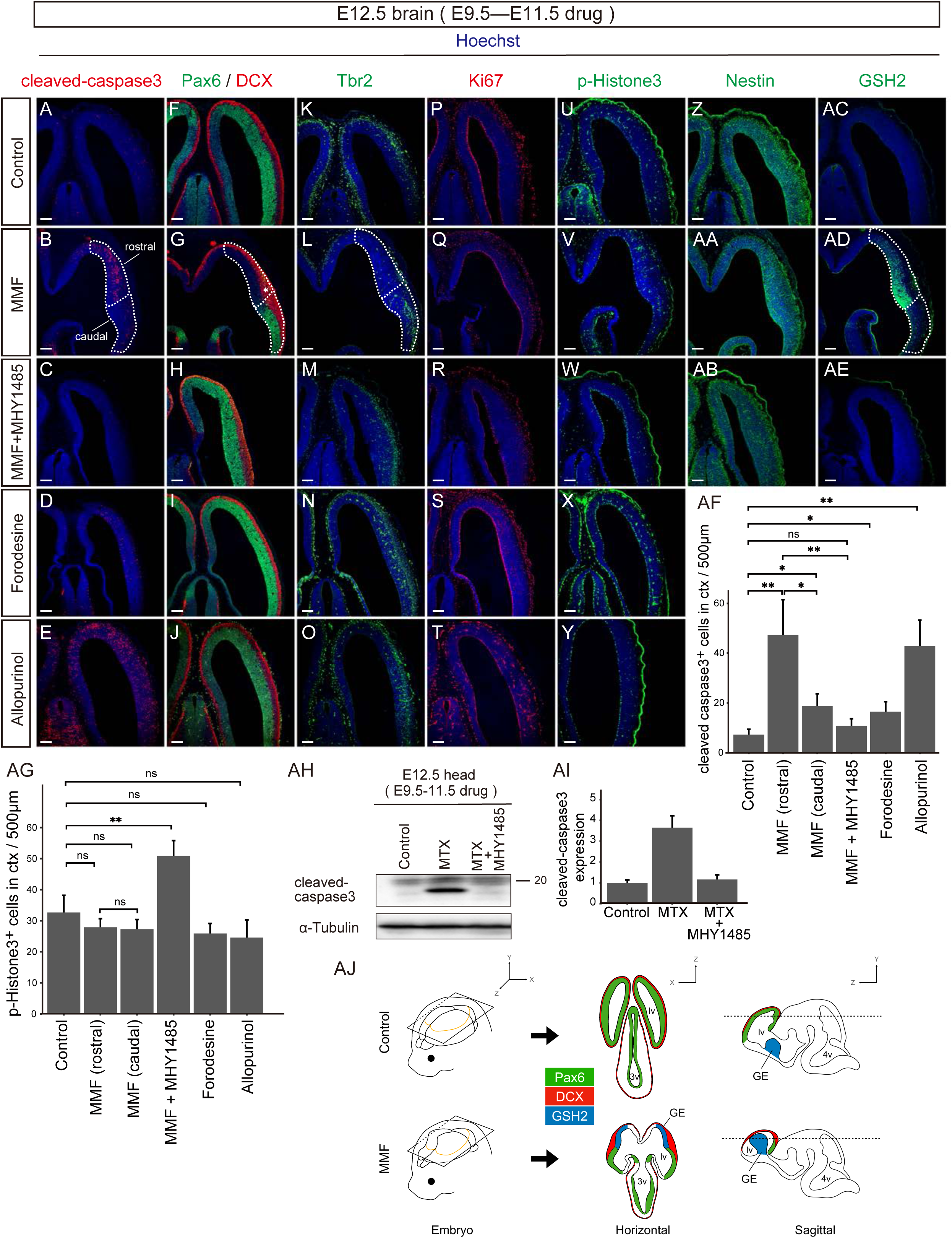
*De novo* pathway inhibition causes forebrain-specific malformation. (A–AE) Control DMSO (A, F, K, P, U, Z, AC), MMF (B, G, L, Q, V, AA, AD), MMF and MHY1485 (C, H, M, R, W, AB, AE), forodesine (D, I, N, S, X), or allopurinol (E, J, O, T, Y) was administered to pregnant mice between E9.5–11.5 and analyzed at E12.5. Horizontal frozen sections were immunostained with antibodies to cleaved-caspase3 (*red*) (A–E), Pax6 (*green*) and DCX (*red*) (F–J), Tbr2 (*green*) (K–O), Ki67 (*red*) (P–T), pH3 (*green*) (U–Y), Nestin (*green*) (Z–AB), or GSH2 (*green*) (AC–AE). The forebrain region treated with MMF was divided by dotted lines into the rostral (Pax6^−^ GSH2^+^) and caudal area (Pax6^+^ GSH2^−^). The asterisk in (G) denotes the expanded cortical layer filled with DCX^+^ immature neurons. (AF, AG) Quantification of cleaved-caspase3^+^ (AF) or pH3^+^ (AG) cells in the cerebral cortex (number of cells per 500 µm width of the cortex). In embryos treated with MMF, cells were counted individually in the rostral and caudal areas. ns, not significant, *p < 0.05, **p < 0.01, Welch’s *t*-test followed by Holm–Bonferroni correction. Scale bars, 100 µm in (A–AE). (AH, AI) Expression of cleaved-caspase3. Embryos were treated with MTX or MTX and MHY1485 during E9.5–E11.5, and the brains were subjected to western blot analysis at E12.5. (AH) Representative western blot illustrating the accelerated apoptosis in MTX-treated brains and the rescue effect of MHY1485. The blot was reprobed with α-Tubulin antibody (bottom) to examine quantitative protein loading. (AI) Quantified comparison of cleaved-caspase3 expression. Data are presented as the mean ± SEM. (AJ) Schematic model of brain abnormalities caused by *de novo* pathway inhibition. Control (left upper panel) or MMF-treated brains (left lower panel) were cut in the horizontal (middle panels) or sagittal plane (right panels). Horizontal sections were sliced through the orange line shown in the 3D brain (left). MMF-treated brains show a loss or hypoplasia of the rostral neocortex and a dorsal expansion of striatal GE (*blue*), resulting in abnormal forebrain structures. In MMF-treated embryos, the rostral neocortex containing Pax6+ VZ disappeared; instead, the GE containing GSH2^+^ VZ ectopically appeared on the dorsal surface. lv, lateral ventricles; 3v, 3rd ventricle; 4v, 4th ventricle; GE, ganglionic eminence.

Next, we performed double immunostaining with anti-Pax6 and anti-DCX, specific markers for RGCs in the cerebral cortex and postmitotic newborn neurons, respectively. The embryonic forebrain typically contains a VZ occupied by Pax6^+^ NSPCs and an IZ/CP occupied by DCX^+^ neurons, as shown in the DMSO-treated control. The structure of the embryonic forebrain treated with forodesine, allopurinol, or MHY1485 was equivalent to that of the control brain, and the spatial distribution of the two cell populations (Pax6^+^ NSPCs and DCX^+^ neurons) appeared unchanged (Fig. 10F, I, J, Extended Data Fig. 10-1). In contrast, in the forebrain of MMF-treated embryos, the distribution of Pax6- and DCX-expressing cells was disrupted and drastically affected: Pax6 expression remained in the caudal area of the forebrain but was completely absent in the rostral and dorsal parts (Fig.10G). With respect to DCX^+^ neurons, although their distribution was relatively preserved throughout the forebrain, the DCX^+^ cortical layer was significantly expanded and thickened at the boundary between the caudal and rostral area (Fig.10G, asterisk). Furthermore, the expression of T-box brain protein 2 (Tbr2) (Englund et al., 2005), an IP marker in the SVZ, was lost in the rostral area of MMF-treated embryos, similar to Pax6 (Fig.10K–O). Nonetheless, the number and distribution of Ki67^+^ or pH3^+^ proliferating cells in rostral area were unaffected (Fig.10P–Y, AG) (control, 32.7 ± 3.0, n = 4; MMF rostral area, 27.9 ± 1.6, n = 8; MMF caudal area, 27.3 ± 1.8, n = 8; forodesine, 25.9 ± 1.8, n = 6; allopurinol, 24.6 ± 3.1, n = 9). Interestingly, co-injection of MMF and MHY1485 increased the number of pH3^+^ proliferating cells in VZ (Fig. 10W, AG) (50.9 ± 4.9, n = 9). Immunostaining with Nestin, a universal RGC marker in all brain regions, unexpectedly revealed that Nestin^+^ RGCs were uniformly distributed in rostral and caudal areas in MMF-treated brains (Fig. 10Z–AB, Extended Data Fig. 10-1F). Since the rostral region of the MTX-treated cortex was occupied by Nestin^+^ but Pax6^−^ RGCs, we considered that this region corresponds to a non-neocortical area region that contains a different subtype of NSPCs.

Previous studies have shown that Pax6 and Tbr2 are expressed in NSPCs of the embryonic cerebral neocortex, while GSH2, another homeobox gene, is expressed explicitly in the NSPC population residing in lateral GE, representing the striatal primordium (Englund et al., 2005; Yun et al., 2001; Corbin et al., 2003). In fact, immunostaining MMF-treated brains with an anti-GSH2 antibody revealed that GSH2 was exclusively expressed in the rostral area, suggesting that this area is a structure derived from the lateral GE (Fig.10AC, AD). Importantly, this phenotype of mislocalization of GSH-positive cells was completely reverted to an intact cortical architecture upon co-treatment with MHY1485 (Fig. 10AE).

Consolidating these histological observations, we concluded that loss or hypoplasia of the rostral neocortex in MMF-treated brains may have caused dorsal displacement and bulging of ventral forebrain regions, including the striatal GE, resulting in abnormal forebrain-like structures (Fig. 10AJ). Consistent with the results in cultured NSPCs, these findings strongly suggest that the NSPC population is more susceptible to deficits in the purine *de novo* pathway regulating mTORC1 signaling pathway in dorsal forebrain regions, such as the neocortex, than in other brain regions. The large number of caspase3^+^ apoptotic cells in the GE-derived region of the rostral area (Fig. 10B, C) also might reflect a shift in dependence on the *de novo* and mTORC1 pathways along the anteroposterior and dorsoventral brain axis.

## Discussion

### Purine synthesis pathways are activated during brain development

We revealed that PAICS and FGAMS, two *de novo* purine synthesis enzymes, were abundant in the embryonic cerebral cortex and downregulated toward postnatal and adult stages. On the other hand, HGPRT, which promotes the salvage pathway, exhibited the opposite trend during cortical development. These results indicate a switch in the main pathway of purine synthesis from the *de novo* pathway to the salvage pathway during brain development. This switch probably reflects that the *de novo* pathway is driven to meet the massive purine demand caused by active cell proliferation during the embryonic period and that the salvage pathway becomes dominant after birth, when the energy cost stabilizes after the peak of neurogenesis and reduced demand for intracellular purines.

The inhibition of *de novo* purine synthesis by drugs *in vivo* caused proliferation defects in NSPCs and delayed the migration of immature neurons in the embryonic neocortex. Previous studies have indicated that purinergic signaling is essential for NSPC maintenance and neuronal migration in the neocortical SVZ during brain development (Lin et al., 2007; Liu et al., 2008; Yamada et al., 2020). Cell cycle progression is closely associated with the properties of NSPCs. For instance, NSPCs residing in the embryonic VZ often exhibit a cellular behavior known as interkinetic nuclear migration (INM), which refers to the periodic movement of the cell nucleus depending on the cell cycle phase. Although the nuclei of NSPCs occupy different positions along the apical/basal axis within the cerebral wall, mitosis (M-phase) exclusively occurs when the nucleus approaches the ventricular surface (apical side) (Baye & Link, 2008; Taverna & Huttner, 2010). Dysregulation of INM is reported to induce the aberrant division of NSPCs on the basal side of the VZ (Tamai et al., 2007). Similarly, upon inhibition of *de novo* synthesis, we observed an abnormal increase in the number of pH3^+^ M-phase cells that lie beyond the VZ (Fig. 6G). The *de novo* purine pathway might play an essential role in INM by controlling cell cycle progression. A premature exit from the cell cycle caused by the inhibition of *de novo* synthesis might delay the radial migration of newborn neurons during brain development.

Additionally, the elaboration of the vascular system within the embryonic cerebral cortex might regulate the demand level for *de novo* purines in NSPCs. Previous studies have indicated the presence of highly vascularized and avascular regions in the developing neocortex and that the nervous system and the vasculature of the brain are closely associated during the development of the cerebral cortex (Komabayashi-Suzuki et al., 2019). An avascular region forms on the VZ from E12.5 to E17.5, where dividing undifferentiated RGCs remain in contact with sprouting neovascular tip cells, which gradually shrinks over development (Bjornsson et al., 2015; Komabayashi-Suzuki et al., 2019). This avascular hypoxic niche is supposedly required to maintain the stemness of NSPCs as well as stem cells in other developing organs (Bjornsson et al., 2015; Komabayashi-Suzuki et al., 2019). As the embryonic NSPCs in the VZ avascular region cannot receive sufficient purines or purine metabolites through blood vessels, it seems logical that NSPCs are strongly dependent on the *de novo* pathway during cortical development. Meanwhile, HGPRT is expressed in CD31^+^ blood vessels during angiogenesis at embryonic stages; a colocalization is not observed at the adult stage (Fig. 3S, Extended Data Fig. 3-1). Thus, the salvage pathway in vascular endothelial cells may supply purines to the surrounding differentiating and migrating neurons in the IZ/CP. In addition, we expected that glial cells, especially astrocytes, which increase in number from late embryonic stages through cell division, would utilize the salvage pathway. However, HGPRT was rarely detected in astrocytes (Fig. 2E). As a recent study revealed that hypoxanthine, an intermediate metabolite of the purine salvage pathway, is involved in morphological changes in microglia (Okajima et al., 2020), microglia might represent the major cell type that actively utilizes the salvage pathway from late embryonic stages to the postnatal stage.

Unlike the embryonic cerebral cortex, the developing cerebellum displayed abundant expression of the salvage enzyme HGPRT in addition to PAICS during the early postnatal period (Fig. 2B, C). Both *de novo* and salvage inhibitors affected the proliferation of cerebellar granular cells, indicating an essential role of both pathways in cerebellar development. One possible reason is that the *de novo* pathway alone may not be sufficient to supply purines since cerebellar NSPCs require large amounts of purines to undergo numerous symmetric cell divisions. Another possible reason is that the cerebellar NSPCs, located in the EGL, could drive the salvage pathway by receiving a supply of purine metabolites from the blood vessels of the meninges covering the brain surface. Over the past decade, a link between purine synthesis pathways and various cancers has been demonstrated (Di Virgilio & Adinolfi, 2017; Yin et al., 2018). In general, the intracellular concentration of purine metabolites and the activity of *de novo* pathway enzymes are enhanced in cancer cells. Changes in the ratio of purine metabolites in tumor cells affect tumor growth, invasion, and metastasis. A switch in purine synthesis pathways that depends on the extracellular environment may not only exist in NSPCs but also in cancer stem cells and other tissue stem cells. This possibility could provide a valuable perspective for anticancer drug development.

### Different purine synthesis pathway dependence of NSPCs according to brain regions

The immunohistochemical analysis of adult mouse brains indicated different dominant purine synthesis pathways according to brain region (Extended Data Fig. 3-2, 3-3). In addition, this study showed that NSPCs have a different predominance of purine synthesis pathways depending on the brain region. Accordingly, disturbance or loss of Pax6, Tbr2, and DCX expression in the neocortex was accompanied by a cortical malformation in which GSH2^+^ GE cells emerged in the dorsal forebrain region. The purine demand in NSPCs likely differs between the dorsal (cerebral neocortex) and ventral forebrain (GE). Considering that NSPCs in the neocortex and GE produce predominantly glutamatergic excitatory and GABAergic inhibitory neurons, respectively, the different cell lineage of each NSPC may explain the differences in the requirement of *de novo* purine synthesis. Indeed, we observed that cell death occurred widely in cells, including neurons in the CP in the rostral GE-like structures formed in MMF-treated embryos (Fig. 10B), suggesting that the cells in GE-like structures belong to distinct cell lineages from their surrounding structures. Alternatively, mitotic delay in NSPCs alters fate specification or viability, as previously described (Pilaz et al., 2016). However, it remains unclear why *de novo* purine production is higher in the neocortex than in other brain regions. There may exist a relationship between purine synthesis pathways and the evolution of mammals with a huge neocortex.

### The *de novo* purine synthesis is involved in cortical formation through mTOR signaling

Our results indicated that inhibiting *de novo* purine synthesis by MMF or MTX during corticogenesis *in vivo* results in brain malformation. Clinical case reports indicate that prenatal exposure to MTX leads to fetal death or fetal MTX syndrome, which is characterized by CNS anomalies, including alobar holoprosencephaly (Corona-Rivera et al., 2010; Seidahmed et al., 2006). However, the molecular mechanism underlying the brain abnormalities induced by MTX remains unknown. We revealed that the brain malformation caused by inhibiting *de novo* purine synthesis was accompanied with a marked inhibition of mTORC1/S6K/S6 signaling and partial impairment of mTORC1/4E-BP1/eIF4E signaling. This malformation was rescued by activation of mTOR signaling. Our results are consistent with previous studies showing that inhibition of purine synthesis in tumor cells mainly suppresses mTORC1/S6K/S6 signaling (Emmanuel et al., 2017; Hoxhaj et al., 2017).

Some patients with cortical malformations, such as hemimegalencephaly and focal cortical dysplasia, have been shown to carry mutations in *mTOR* (Tarkowski et al., 2019). Deficiency in *mTOR* causes morphological abnormalities in the brain, including a thinner cerebral cortex due to suppressed NSPC proliferation (Ka et al., 2014), suggesting a close correlation between the mTOR pathway and disruption of cortical development. These findings further supported our notion that *de novo* purine synthesis and the mTORC1/S6K/S6 signaling pathway tightly regulate each other spatiotemporally to control neocortical development.

### Limitations

In this study, we revealed a switch from *de novo* to salvage purine synthesis as brain development proceeds, and the *de novo* pathway plays a vital role in early embryonic corticogenesis, cooperating with the mTOR signaling pathway. We indicated that purine sensibility or predominance of purine synthesis pathways differs in NSPCs depending on the brain region. However, we could not clarify the factor that drives the switch from *de novo* to salvage pathways as well as the molecular mechanisms of brain malformation caused by *de novo* purine synthesis inhibitors. Our future study will focus on regional and lineage-specific differences in purine metabolism in the brain. For example, genetic manipulation of the *de novo* synthesis enzymes using Emx1-Cre transgenic mice will evidently provide molecular and cellular clues regarding the role of this pathway in determining the properties of NSPCs in the dorsal telencephalon. These studies might shed light on the significance of adaptive changes in purine metabolic pathways during the evolution of the mammalian cerebral neocortex.

## Acknowledgments

We would like to thank enago (www.enago.com) for English language editing.

