## supplemental file for "Spatiotemporal regulation of *de novo* and salvage purine synthesis during brain development"

### Extended Data Legends

**Figure 3-1. HGPRT is expressed in CD31$^{+}$ endothelial cells in the developing cerebral cortex**

E16.5 (A) and E18.5 (B) cerebral cortices double immunostained with anti-CD31 (*green*) and anti-HGPRT (*red*) antibodies. Insets present magnified views of each blood vessel. Scale bar, 50 μm.

**Figure 3-2. Expression of purine synthesis enzymes in the adult brain**

(A–X) Coronal sections of adult brains immunostained with anti-PAICS (A–H), anti-FGAMS (I–P), and anti-HGPRT (Q–X) antibodies. Representative images of the locus coeruleus (LC) (A, I, Q), Edinger–Westphal nucleus (EW) (B, J, R), vestibular nucleus (VN) (C, K, S), medial vestibular nucleus (Med) (D, L), ambiguous nucleus (Amb) (E, M), facial nucleus (7N) (F, N), trigeminal motor nucleus (Ⅴmo) (G), ventral cochlear nucleus (VC) (H), medial habenular nucleus (MHb) (O, X), islands of Calleja (ICj) (P), arcuate hypothalamic nucleus (Arc) (T), zona incerta (ZI) (U), nigrostriatal bundle (NS) (V), and lateral septal (LS) (W). Areas surrounded by a dashed line in (A) denote the LC and mesencephalic trigeminal nucleus (Me5). Scale bar, 20 μm.

**Figure 3-3. Expression of purine synthesis enzymes in the adult brain**

(A–P) Coronal sections of adult brains immunostained with anti-PAICS (A, D, G, J, M), anti-FGAMS (B, E, H, K, N), and anti-HGPRT (C, F, I, L, O) antibodies. Representative images of the cerebellum (cbl) (A–C), lateral cerebellar nucleus (Lat) (D–F), red nucleus (RN) (G–I), posterior interlaminar thalamic (PIL) complex (J–L), and polymorph layer of the dentate gyrus (PoDG) (M–O). (P) Expression profiles of PAICS, FGAMS, and HGPRT in discrete brain regions and nuclei. Black circles indicate regions with high immunoreactivity. ML, molecular cell layer; PCL, Purkinje cell layer; GCL, granule cell layer; GrDG, granule cell layer of the dentate gyrus. Scale bar, 100 µm in (A–O).

**Figure 4-1. Inhibition of *de novo* purine synthesis disturbs the proliferation of cerebellar NSPCs *in vitro***

Primary cultured NSPCs derived from P1 EGL were treated for 48 h with PBS or DMSO (control, A), forodesine (1 and 5 µM) (B), or MMF (1, 5, and 10 µM) (C), followed by BrdU labeling for 24 h. NSPCs were immunostained with anti-Nestin (*green*) and anti-BrdU (*red*) antibodies. (D) Quantification of dividing NSPCs. The ratio of the number of BrdU$^{+}$ Nestin$^{+}$ cells to the total number of Nestin$^{+}$ cells. ns, not significant; ***$p < 0.001$, chi-squared test with Holm–Bonferroni correction. Control PBS, n = 460; control DMSO, n = 396; forodesine 1 µM, n = 373; forodesine 5 µM, n = 700; MMF 1 µM, n = 73; MMF 5 µM, n = 70; MMF 10 µM, n = 64. Scale bar, 50 µm.

**Figure 6-1. Expression of GFAP in cerebral cortices treated with purine synthesis inhibitors**

E12.5 embryos were treated with control DMSO (A), forodesine (B), or MMF (C) and analyzed at E14.5. Horizontal frozen sections were immunostained with anti-GFAP antibody (*red*). Nuclei were stained with Hoechst dye (*blue*). MZ, marginal zone; CP, cortical plate; IZ, intermediate zone; SVZ/VZ, subventricular zone/ventricular zone.

Scale bar, 100 μm in (A–F).

**Figure 7-1. 4E-BP1 expression**

Quantified comparison of the total protein content of 4E-BP1 (A) and each isoform [4E-BP1-α (B), 4E-BP1-β (C), or 4E-BP1-γ (D)] in Figure 7A. Protein bands obtained using three independent mouse brains treated with each drug (DMSO, MTX, allopurinol, or MTX and allopurinol) were quantified using α-tubulin as an internal standard. Data are presented as the mean ± SEM.

**Figure 9-1. Brain malformations caused by the inhibition of the *de novo* pathway**

Serial horizontal sections of each E12.5 embryo were treated with different inhibitors. The left and right panels represent the sections cutting through the dorsal and ventral planes, respectively. The pregnant mice were successively treated with control DMSO (A), forodesine (B), MMF (C), MHY1485 (D), MMF and MHY1485 (E), MTX (F), allopurinol (G), or MTX and allopurinol (H) during E9.5–E11.5, and the brains were harvested at E12.5. Sections were stained with Hoechst dye (cyan). Scale bar, 500 μm.

**Figure 10-1. Embryonic brains treated alone with the mTOR activator MHY1485**

MHY1485 was administered to pregnant mice between E9.5 and E11.5, followed by analysis at E12.5. Horizontal frozen sections were immunostained with antibodies against cleaved-caspase3 (*red*) (A), Pax6 (*green*)/DCX (*red*) (B), Tbr2 (*green*) (C), Ki67 (*red*) (D), pH3 (*green*) (E), Nestin (*green*) (F), or GSH2 (*green*) (G). Scale bar, 100 μm in (A–G).

Fig.3-1

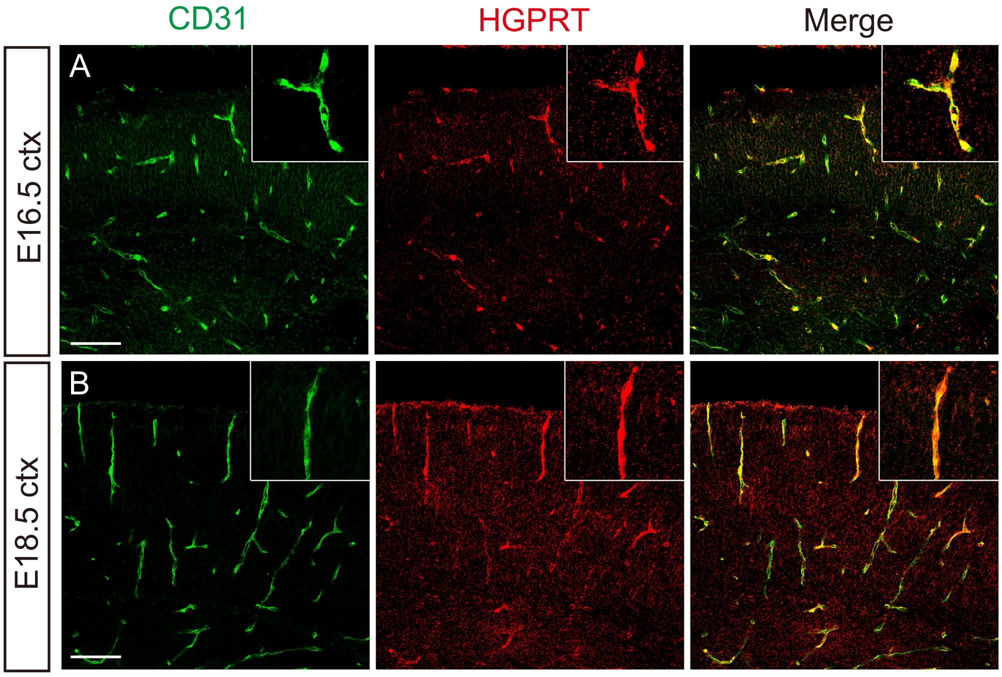

Fig.3-2

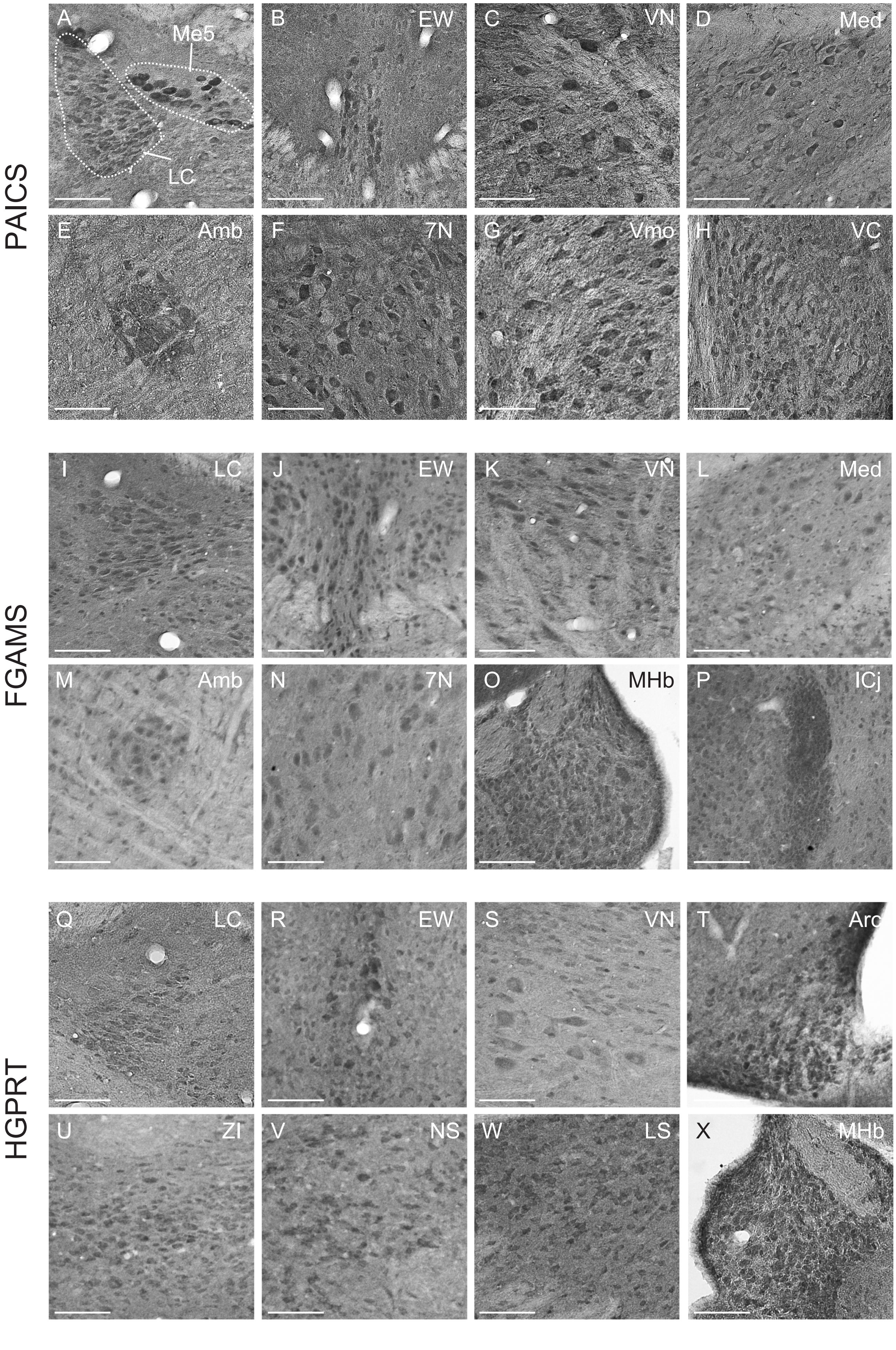

PAICS
A
Me5
LC
B
EW
C
VN
D
Med
E
Amb
F
7N
G
Vmo
H
VC
FGAMS
I
LC
J
EW
K
VN
L
Med
M
Amb
N
7N
O
MHb
P
ICj
HGPRT
Q
LC
R
EW
S
VN
T
Arc
U
ZI
V
NS
W
LS
X
MHb

Fig.3-3

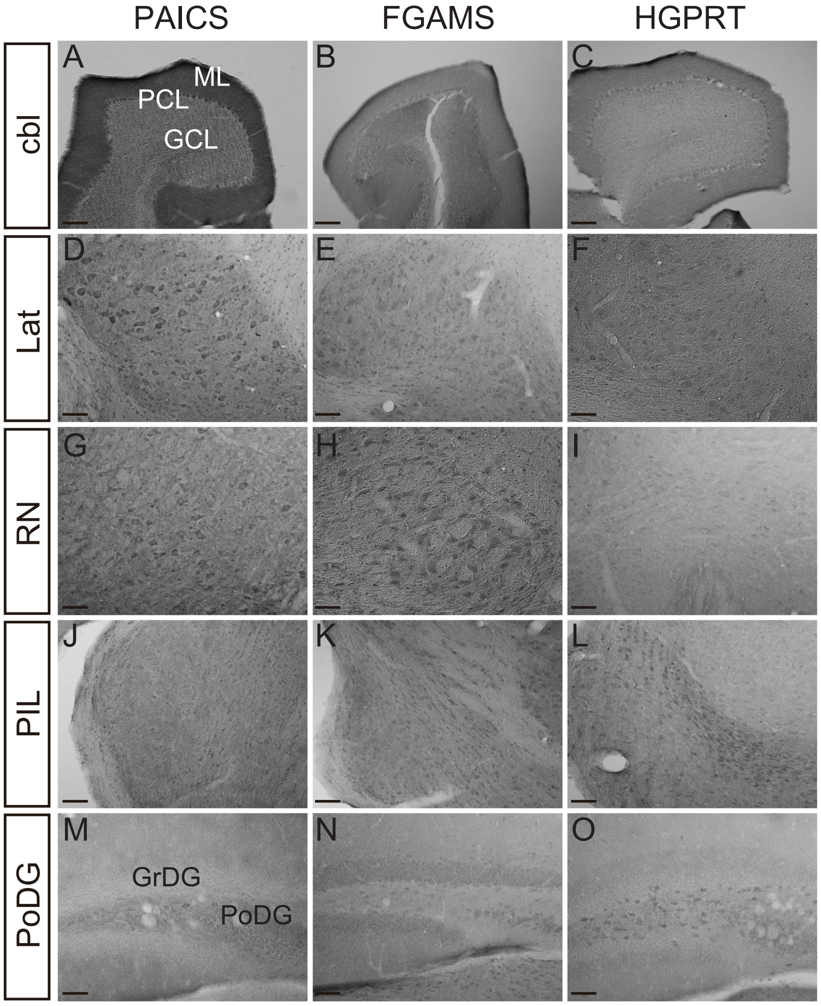


P

| Brain area | PAICS | FGAMS | HGPRT |
|---|---|---|---|
| locus coeruleus (LC) | ● | ● | ● |
| Edinger-Westphal nucleus (EW) | ● | ● | ● |
| vestibular nucleus (VN) | ● | ● | ● |
| lateral cerebellar nucleus (Lat) | ● | ● | |
| red nucleus (RN) | ● | ● | |
| facial nucleus (7N) | ● | ● | |
| medial vestibular nucleus (Med) | ● | ● | |
| ambiguous nucleus (Amb) | ● | ● | |
| motor trigeminal nucleus (Vmo) | ● | | |
| mesencephalic trigeminal nucleus (Me5) | ● | | |
| pontine reticular nucleus (PnC) | ● | | |
| trochlear nucleus (4N) | ● | | |
| ventral cochlear nucleus (VC) | ● | | |
| medial habenular nucleus (MHb) | | ● | ● |
| islands of calleja (ICj) | | ● | |
| posterior interlaminar thalamic (PIL) | | | ● |
| polymorph layer of dentate gyrus (PoDG) | | | ● |
| arcuate hypothalamic nucleus (Arc) | | | ● |
| dorsomedial hypothalamic nucleus (DMN) | | | ● |
| zona incerta (ZI) | | | ● |
| lateral septal (LS) | | | ● |
| nigrostriatal bundle (NS) | | | ● |

Fig.4-1

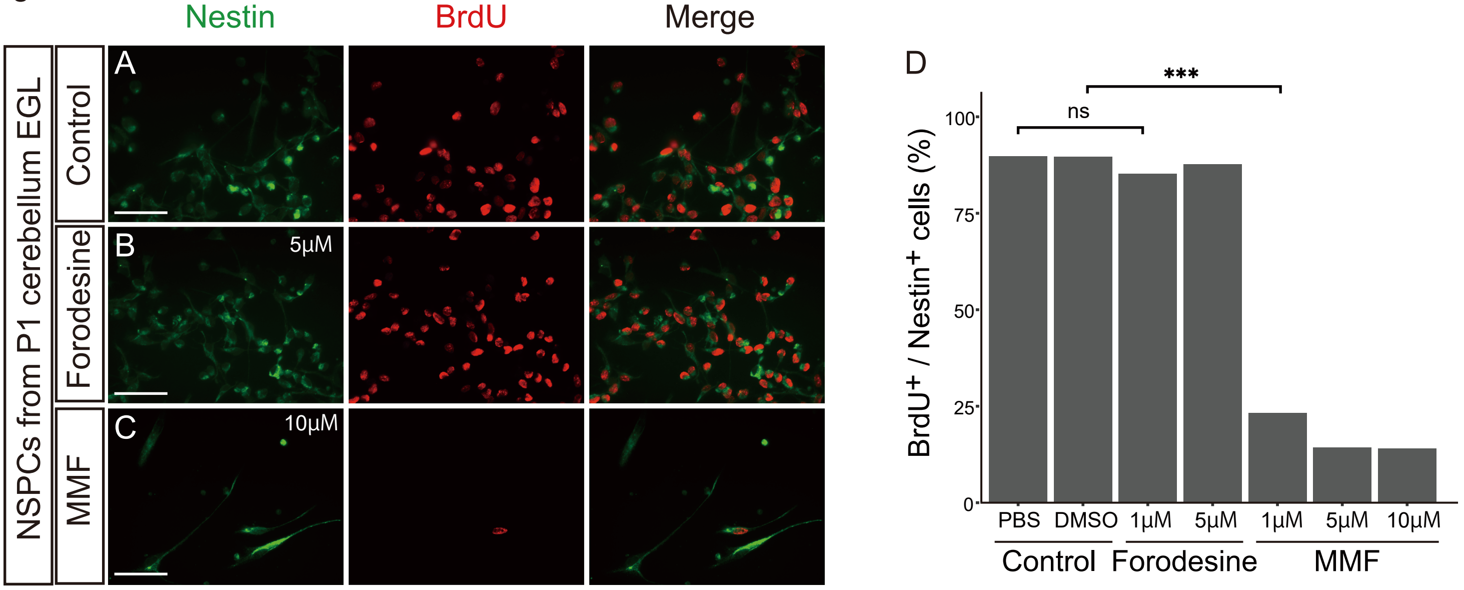
Nestin
BrdU
Merge
NSPCs from P1 cerebellum EGL
Control
Forodesine
MMF
A
B
5μM
C
10μM
D
BrdU+ / Nestin+ cells (%)
100
75
50
25
0
ns
***
PBS
DMSO
1μM
5μM
1μM
5μM
10μM
Control
Forodesine
MMF

Fig.6-1

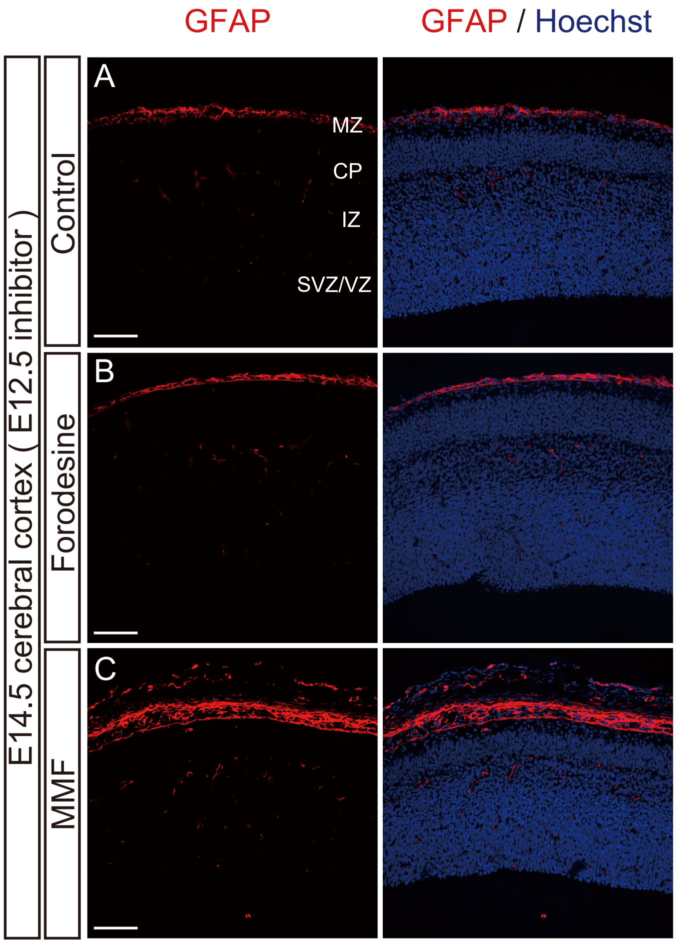
GFAP
GFAP / Hoechst
E14.5 cerebral cortex ( E12.5 inhibitor )
Control
Forodesine
MMF
A
B
C
MZ
CP
IZ
SVZ/VZ

Fig.7-1

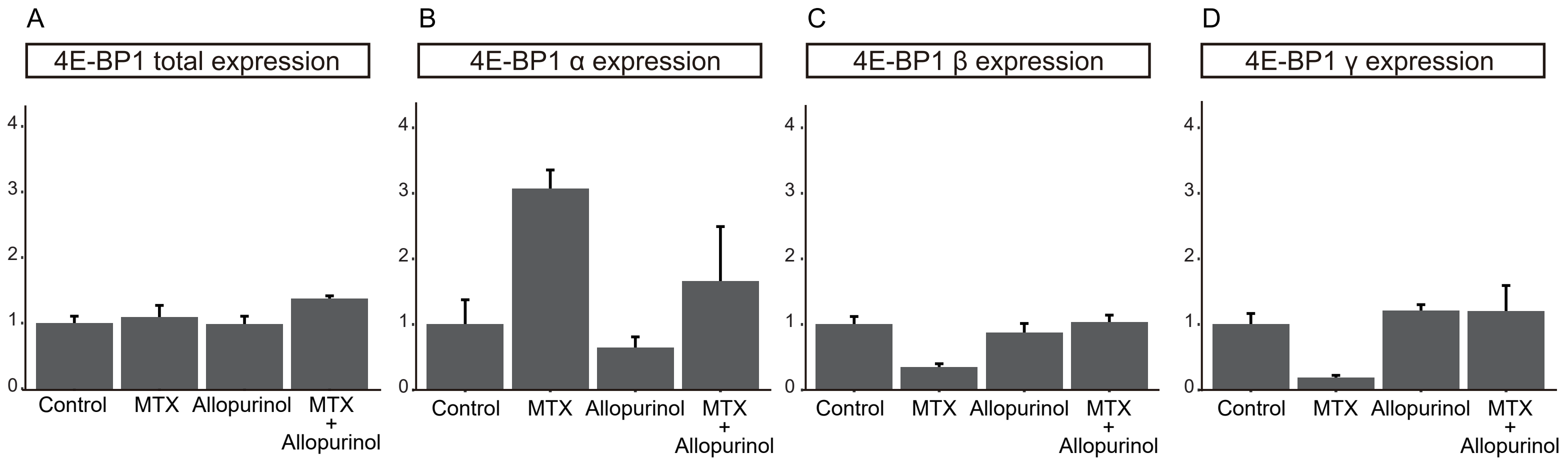
A
4E-BP1 total expression
B
4E-BP1 α expression
C
4E-BP1 β expression
D
4E-BP1 γ expression
0
1
2
3
4
Control
MTX
Allopurinol
MTX
+
Allopurinol

Fig.9-1

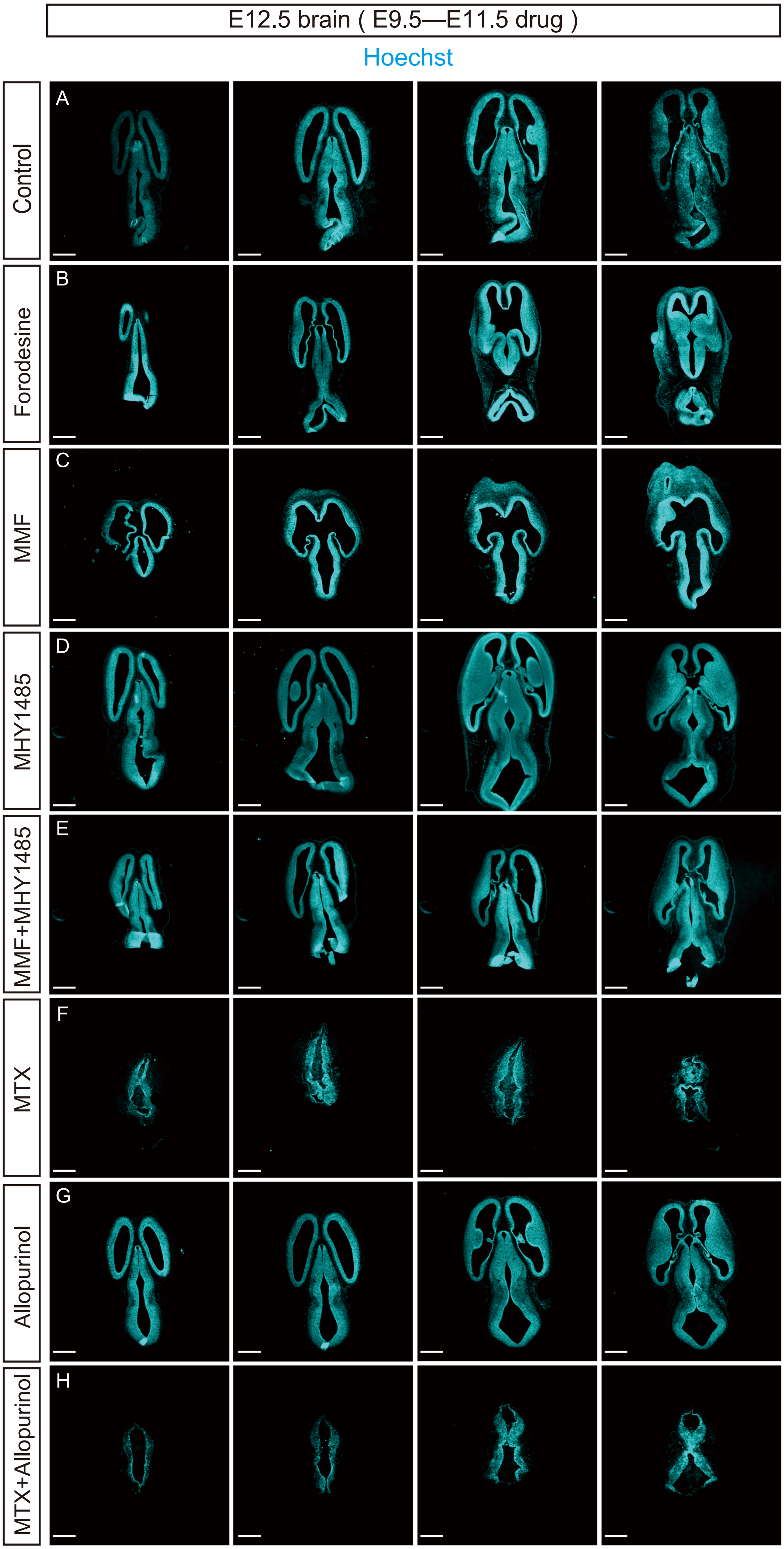

E12.5 brain ( E9.5—E11.5 drug )
Hoechst
Control
A
Forodesine
B
MMF
C
MHY1485
D
MMF+MHY1485
E
MTX
F
Allopurinol
G
MTX+Allopurinol
H

Fig.10-1

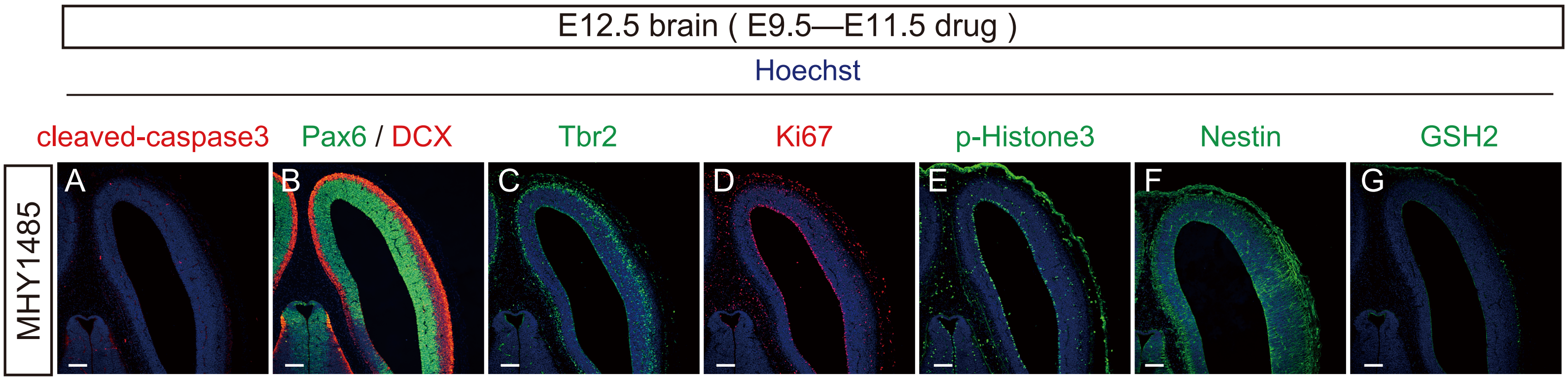

E12.5 brain ( E9.5—E11.5 drug )
Hoechst
cleaved-caspase3
Pax6 / DCX
Tbr2
Ki67
p-Histone3
Nestin
GSH2
MHY1485
A
B
C
D
E
F
G

Table1

**One way ANOVA followed by Welch's t test with Holm–Bonferroni correction**

| Fig. 5E | T | 95.00% CI of difference | Adjusted *p* value | Summary |
|---|---|---|---|---|
| Control vs MMF | -3.245 | -22.99 to -4.511 | 0.007082 | ** |
| Control vs Forodesine | -4.801 | -28.51 to -10.81 | 0.0006982 | *** |
| Control vs MTX | -6.532 | -38.48 to -19.45 | 0.00003999 | *** |

| Fig. 5I | T | 95.00% CI of difference | Adjusted *p* value | Summary |
|---|---|---|---|---|
| Control vs MMF | -4.746 | -22.83 to -8.520 | 0.0008252 | *** |
| Control vs Forodesine | -2.932 | -19.94 to -2.504 | 0.01749 | * |

| Fig. 6D | T | 95.00% CI of difference | Adjusted *p* value | Summary |
|---|---|---|---|---|
| Control (cp) vs MMF (cp) | 7.139 | 5.472 to 10.68 | 0.0004716 | *** |
| Control (cp) vs Forodesine (cp) | 4.027 | 2.666 to 8.680 | 0.004504 | ** |
| Control (iz) vs MMF (iz) | 3.781 | 6.492 to 23.87 | 0.007065 | ** |
| Control (iz) vs Forodesine (iz) | 0.6651 | -5.523 to 10.47 | 0.5171 | ns |
| Control (svz/vz) vs MMF (svz/vz) | -6.261 | -31.52 to -15.00 | 0.0005173 | *** |
| Control (svz/vz) vs Forodesine (svz/vz) | -2.417 | -15.59 -0.7031 | 0.0695 | ns |

| Fig. 6H | T | 95.00% CI of difference | Adjusted *p* value | Summary |
|---|---|---|---|---|
| Control vs MMF | 3.594 | 5.616 to 22.57 | 0.006586 | ** |
| Control vs Forodesine | -0.2730 | -9.961 to 7.699 | 0.7886 | ns |

| Fig. 7B | T | 95.00% CI of difference | Adjusted *p* value | Summary |
|---|---|---|---|---|
| Control vs MTX | 19.90 | 0.7197 to 1.101 | 0.008576 | ** |
| Control vs Allo | 0.4232 | -0.2907 to 0.3780 | 0.7015 | ns |
| Control vs MTX + Allo | 0.5208 | -1.209 to 1.555 | 1 | ns |

| Fig. 10AF | T | 95.00% CI of difference | Adjusted *p* value | Summary |
|---|---|---|---|---|
| Control vs MMF (rostral) | -4.844 | -59.06 to -21.16 | 0.00675 | ** |
| Control vs MMF (caudal) | -3.735 | -18.36 to -4.749 | 0.01310 | * |
| Control vs MMF + MHY1485 | -1.716 | -7.973 0.8622 | 0.1068 | ns |
| Control vs Forodesine | -3.466 | -15.01 to -3.437 | 0.01362 | * |
| Control vs Allopurinol | -5.841 | -49.55 to -21.78 | 0.001961 | ** |
| MMF (rostral) vs MMF (caudal) | 3.297 | 9.229 to 47.88 | 0.01635 | * |
| MMF (rostral) vs MMF + MHY1485 | 4.378 | 17.55 to 55.56 | 0.00977 | ** |

| Fig. 10AG | T | 95.00% CI of difference | Adjusted *p* value | Summary |
|---|---|---|---|---|
| Control vs MMF (rostral) | 1.323 | -3.087 to 12.67 | 0.8404 | ns |
| Control vs MMF (caudal) | 1.456 | -2.626 to 13.46 | 0.8460 | ns |
| Control vs MMF + MHY1485 | -4.264 | -27.29 to -9.153 | 0.003657 | ** |
| Control vs Forodesine | 1.833 | -1.216 to 14.77 | 0.2699 | ns |
| Control vs Allopurinol | 1.770 | -1.605 to 17.83 | 0.1916 | ns |
| MMF (rostral) vs MMF (caudal) | 0.2430 | -4.897 to 6.147 | 0.8116 | ns |

**Chi-square test with Holm–Bonferroni correction**

| Fig. 4D | χ2 | Adjusted *p* value | Summary |
|---|---|---|---|
| Control DMSO vs MMF 1μM | 145.5 | 3.33353E-33 | *** |
| Control PBS vs Forodesine 0.1μM | 0.03718 | 0.4002 | ns |

| Fig. 4G | χ2 | Adjusted *p* value | Summary |
|---|---|---|---|
| Control shRNA vs PAICS shRNA #1 | 18.03 | 2.176E-05 | *** |
| Control shRNA vs PAICS shRNA #3 | 80.27 | 6.526E-19 | *** |

| Fig. 4-1D | χ2 | Adjusted *p* value | Summary |
|---|---|---|---|
| Control DMSO vs MMF 1μM | 165.5 | 1.457E-37 | *** |
| Control PBS vs Forodesine 1μM | 1.170 | 0.2794 | ns |

| Fig. 8E | χ2 | Adjusted *p* value | Summary |
|---|---|---|---|
| Control 6h vs MMF 6h | 33.67 | 1.956E-08 | *** |
| Control 6h vs Forodesine 6h | 1.247 | 0.2641 | ns |
| MMF 6h vs MMF+MHY1485 6h | 13.56 | 0.0004622 | *** |
| Control 48h vs MMF 48h | 18.21 | 5.933E-05 | *** |
| Control 48h vs Forodesine 48h | 0.2464 | 0.6196 | ns |
| MMF 48h vs MMF+MHY1485 48h | 6.956 | 0.01671 | * |

| Fig. 8J | χ2 | Adjusted *p* value | Summary |
|---|---|---|---|
| Control vs MMF | 0.03718 | 0.8471 | ns |
| Control vs Forodesine | 1.528 | 0.4329 | ns |
